# An aged human heart tissue model showing age-related molecular and functional deterioration resembling the native heart

**DOI:** 10.1101/287334

**Authors:** Aylin Acun, Trung Dung Nguyen, Pinar Zorlutuna

**Affiliations:** Bioengineering Graduate Program, University of Notre Dame, IN; Aerospace and Mechanical Engineering Department, University of Notre Dame, IN

**Keywords:** aging, tissue engineering, reperfusion injury, human induced pluripotent stem cell

## Abstract

Deaths attributed to ischemic heart disease increased by 41.7% from 1990 to 2013. This is primarily due to an increase in the aged population, however, research on cardiovascular disease (CVD) has been overlooking aging, a well-documented contributor to CVD. The field heavily depends on the use of young animals due to lower costs and ready availability, despite the prominent differences between young and aged heart structure and function. Here we present the first human induced pluripotent stem cell (hiPSC)-derived cardiomyocyte (iCM)-based, in vitro aged myocardial tissue model as an alternative research platform. Within 4 months, iCMs go through accelerated senescence and show cellular characteristics of aging. Furthermore, the model tissues fabricated using these aged iCMs, with stiffness resembling that of aged human heart, show functional and pharmacological deterioration specific to aged myocardium. Our novel tissue model with age-appropriate physiology and pathology presents a promising new platform for investigating CVD or other age-related diseases.

## Introduction

Cardiovascular disease (CVD) is the leading global cause of death since 1910^1^. The fact that CVD remains the number one killer for over a century suggests that, despite the advancements in science and medicine over the years, there is a huge gap in translating these scientific findings to clinical setting. One of the major reasons for this gap is pre-clinical research’s heavy dependence on young animal models despite the fact that aging is the biggest risk factor for CVD. The average age for first heart attack is 65.3 years for males and 71.8 years for females^1^. As such, understanding and implementing the age-related phenotype of the heart tissue is crucial for studying CVD in general, and myocardial infarction (MI) specifically. Here we present an alternative model that is of human-origin and that mimics aging on cellular and tissue level as we have studied in detail through characterization and functional validation of this novel tissue model in comparison with aged human heart tissue.

Aging affects the structure and functionality of the heart cells as well as the extracellular matrix (ECM) surrounding them. The well-documented changes at organ level include the thickening and stiffening of the heart wall. Importantly, the cardiac reserve is lowered and the effectiveness of the repair system declines, rendering heart more vulnerable to disease and stress^2^. In addition, it was shown that the stiffness of individual cardiomyocytes also increase with age^3^. It was also reported that there is an approximately 35% decrease in the total number of ventricular cardiomyocytes^4^ leading to enlargement of remaining cells^5^. Functionally, aging in the heart is associated with prolonged contraction and relaxation times of cardiomyocytes^6,7^ as well as with an age-related reduction in responsiveness to β-adrenergic receptor stimulation^8–10^. With advanced age, cardiomyocytes are reported to enter senescence as identified with an increase in p21 and p53 expression levels at later ages^5,11^ and a related lipofuscin granule presence was observed in the aged heart^12^. Pharmacologically, Jamieson et al. have shown that soluble epoxide hydrolase deletion had a protective effect on 3-month-old mice following MI, however, this protective effect was significantly impaired in 16-month-old mice^13^. Similarly, other studies showed that pharmacological effects of various drugs such as adrenaline^14^, isoproterenol, propranolol, and salbutamol^10,15^ were reduced in individuals older than 70, showing the crucial role of aging in heart physiology and pathophysiology.

Currently, despite the prominent differences between young and aged heart structure and function, using young animals to study diseases that abundantly affect the elderly population is preferred due to its lower cost and lack of comparable historical data for aged animals^16–19^. Jackson et al. has reported that the most common rodent age used in the 297 studies conducted was 8-12 weeks. Strikingly, all of the MI studies included in the survey used animals 12 weeks or younger, equivalent to humans 20-25 of age^19^. Therefore, the role of aging is significantly overlooked in many existing animal studies. In addition, there are prominent differences between human and animal physiology and pathology^20^. The recent advancements in tissue engineering now allows for the fabrication of complex, 3 dimensional (3D) engineered tissue models in a highly controlled manner^21,22^. Using the human induced pluripotent stem cells (hiPSCs), these models can potentially be personalized to study drug testing and cytotoxicity assessments on a human-origin platform^23^. The design of such engineered models should take into consideration the cellular and microenvironmental factors related to the disease under investigation. Therefore, a tissue engineered platform incorporated with cells and ECM which show aging characteristics may present a potent alternative to using young animals to study CVDs such as MI, on tissue-level systems.

Here we characterized hiPSC-derived cardiomyocytes (iCMs) over a prolonged culture period and determined that maturation process reaches a limit around 55 days in culture and is followed by a functional deterioration at late stages of culture (after 100 days). We show, for the first time, an in vitro aged, human-origin cardiac tissue model that is both physiologically and pathologically relevant for studying CVD such as MI. We achieved this by incorporating iCMs at the late stages into matrices with aged human heart stiffness. We show that late iCMs at cellular level and the respective tissue models at tissue level behave similar to aged human hearts, showing age-related responses, including altered gene expression and proliferative properties, impaired functional properties and stress response, decreased sensitivity to therapeutics and lower long-term viability.

## Results

### Phenotypical characterization of iCMs with respect to culture age

In the current state of the literature, the relation between the purity and maturity of the iPSC-derived cardiomyocytes with respective to their culture age has not been fully established. Therefore, we first characterized iCM purity and functionality extensively, in relation to the culture age over a 4-month culture period and established a reproducible cell source for fabricating the heart tissue models with different age characteristics. We assessed the phenotypic and biochemical properties of the iCMs on days 17, 21, 35, 55, 75, 100, and 120 of culture to acquire a thorough understanding of their maturity with respect to culture age. To avoid any idiosyncratic behavior coming from one particular donor, or reprogrammed cell type, we differentiated 2 different hiPSC lines from two different sources: a fibroblast-origin cell line (DiPS-1016SevA)^24^, and an <u>e</u>ndothelial-origin cell line (Huv-iPS4F1)^25^, using a previously established protocol^26^, yielding <u>F</u>-iCMs and <u>E</u>-iCMs, respectively. We first characterized the differentiation reproducibility of these two cell lines. Our results showed that both cell lines yielded strongly beating iCMs in 85% of the differentiations (49 batches of differentiation), indicating that both cell lines are suitable for reproducible iCM differentiation (Figure 1A). We also determined the differentiation efficiency of each batch using FACS against cardiac troponin T2 (TNNT2) on day 21 and represented as TNNT2 positive cell percentage within each of the beating populations. Over 90% of the cells were TNNT2 positive, showing that the differentiation is successful with high yield and purity without the need of any post-purification after differentiation completion (Figure 1B). Next, we characterized the change in spontaneous beat rate with culture age (Supplementary videos 1-3). Both cell lines showed the highest spontaneous beat rate between days 35-55 of culture, followed by a decrease with age (Figure 1C). The highest beat rate for F-iCMs was recorded to be 42±8.3 beats per minute on day 55 while the highest beat rate for E-iCMs was recorded to be 41±2.5 beats per minute on day 35 of culture. For both cell lines decline in beat rate started on day 75 and the lowest bpm was reached on day 120 of culture. We also qualitatively characterized the structural maturity of iCMs through TNNT2 immunostaining.

**Figure 1.**
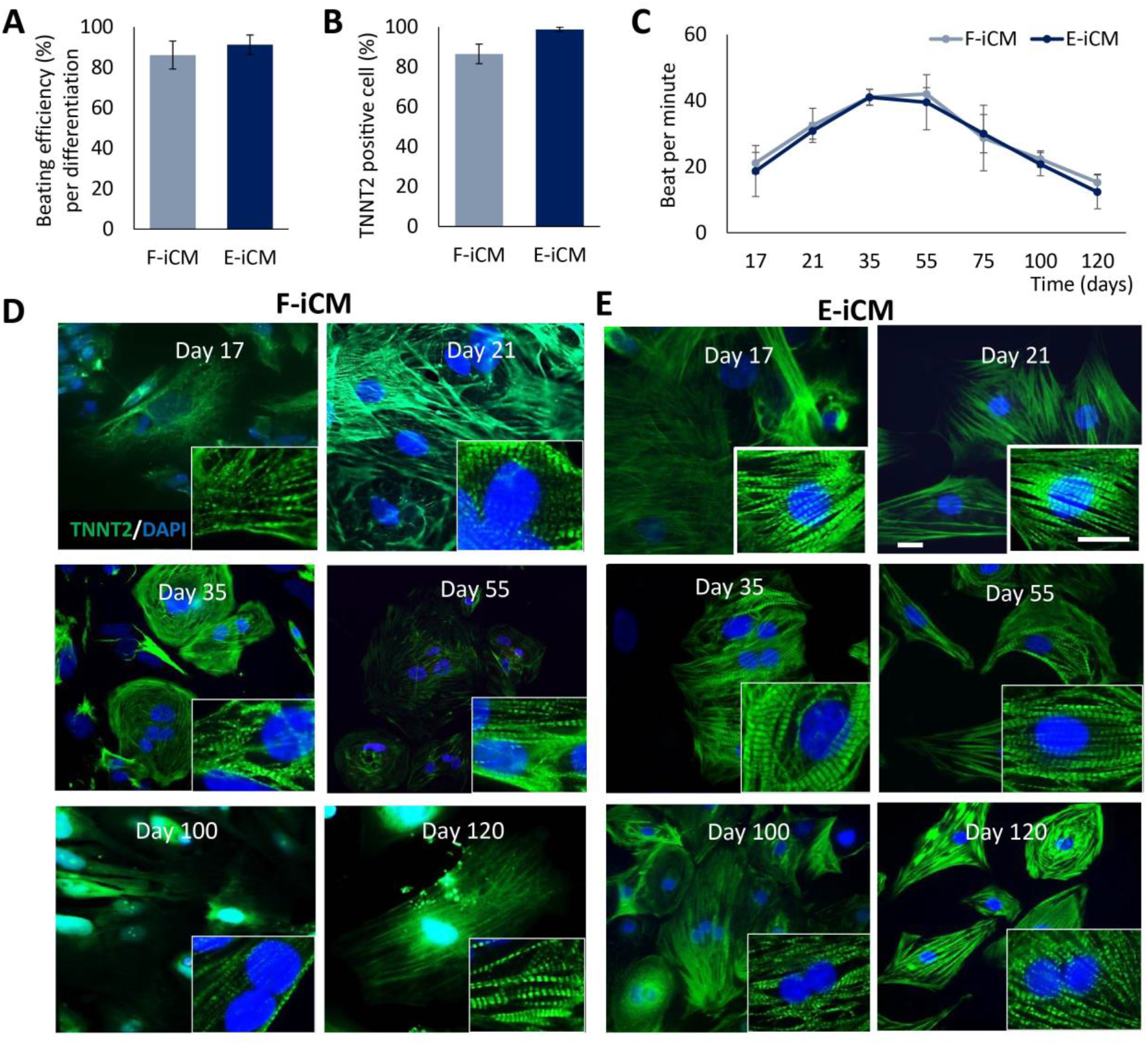
iCM differentiation efficiency characterization. (A) The beating efficiency (%) per differentiation of F-iCMs and E-iCMs. (B) The FACS analysis against TNNT2 of F-iCMs and E-iCMs showing differentiation efficiency of each differentiation batch represented as TNNT2 positive cell (%). (C) The Spontaneous beat rate per E-iCMs at different days of culture. The TNNT immunostaining of (D) F-iCMs, and (E) E-iCMs on different days of culture. Cell nuclei are stained with DAPI (blue). (Scale bars=20 µm) (n≥6.)

Both cell lines showed striated TNNT2 structures, an indicator of healthy cardiomyocyte morphology, from day 17-120 in culture (Figure 1D, E), showing that the differentiation to iCMs was successful and that iCMs do not lose their healthy structure throughout the 120 days in culture.

### Structural and Molecular Maturation of iCMs with Culture Age

We sought to identify the culture age at which iCMs reach molecular and functional maturity throughout the 120 days of culture. We have selected 8 genes based on previous studies where global gene expression in fetal and adult human heart tissue, and human PSC-derived cardiomyocytes were investigated. Seven genes, TNNT2, S100A1, MYOM3, MYL2, MYH6, CASQ2, COX6A2, were shown to be expressed at significantly higher levels in adult hearts, compared to fetal hearts^27–29^. Thereby, the change in expression of these 7 markers indicates the structural (TNNT-2, MYL2, MYH6, MYOM3) and functional (CASQ2, S100A1 are involved in excitation-contraction coupling and COX6A2 is involved in respiration) maturation of iCMs in long term culture. We compared the expression levels of these markers in iCMs to their expression in 65-year-old human heart tissues to find out the relevance of in vitro culture time with maturity of iCMs. We found that expression of TNNT2, S100A1, CASQ2, COX62A, MYOM3, and MYH6 increased significantly after day 21 of culture for both F-iCMs (Figure 2A) and E-iCMs (Figure 2B). MYL2 expression significantly increased with time in both cell lines as well, but this increase was significant after day 35 in F-iCMs. The expression levels stabilized after day 35 or 55 for E-iCMs, and after day 75 for F-iCMs. Importantly, the TNNT2, MYH6, and MYL2 expression of day 75 and day 120 F-iCMs, and TNNT2 and MYL2 expression of day 55-120 E-iCMs were comparable to the expression levels in native human heart tissues. The significantly lower expression of S100A1, CASQ2, MYOM3, and COX6A2 in iCMs, compared to the expression levels in 65-year-old native human tissue is in agreement with other reports of hPSC-derived cardiomyocytes^27–29^. We also monitored NKX2.5 expression, as NKX2.5 is known to be involved in heart development^30^ and has been shown to decrease as the heart matures^31^. The NKX2.5 expression was highest in day 21 F-iCMs and day 21-35 E-iCMs and decreased gradually with culture time reaching the lowest values on day 120 of culture. NKX2.5 expression in iCMs was comparable to the expression levels in native human tissue only after day 55 for both F-iCMs and E-iCMs supporting the trend we observed in maturity marker expression. As expected, the undifferentiated iPSC controls’ expression of all 8 cardiac marker genes was significantly lower than both native human tissue and iCMs.

**Figure 2.**
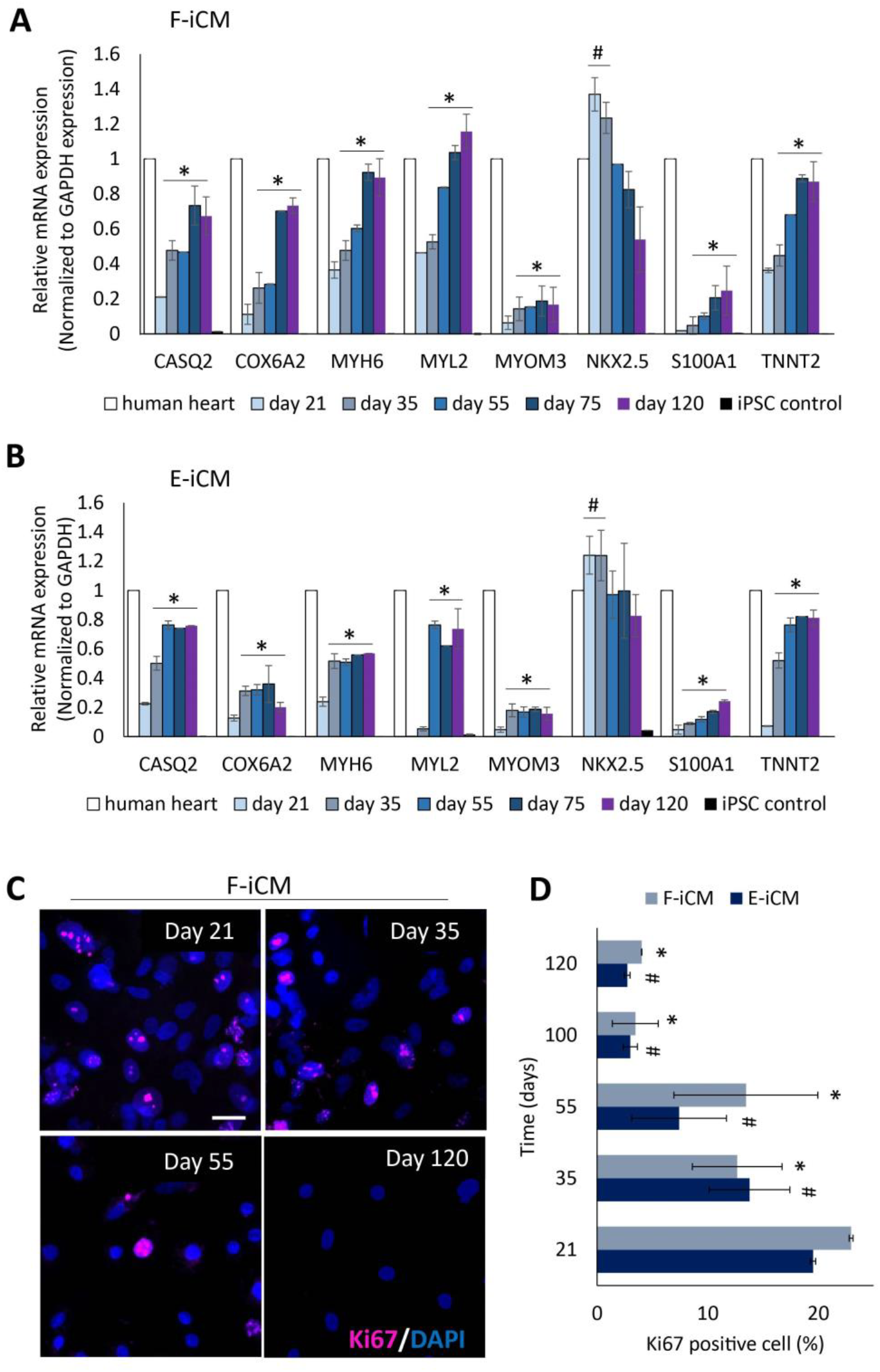
Molecular characterization of iCMs. RT-qPCR analysis of relative mRNA expression of cardiac markers CASQ2, COX6A2, MYH6, MYL2, MYOM3, NKX2.5, S100A1, and TNNT2 in 65-year-old human heart and (A) F-iCMs, and (B) E-iCMs at different culture ages. Expression in human heart was normalized to 1 for easier comparison. * represents p<0.05 significant difference in expression of a single target compared to day 21 (One-way ANOVA followed by Tukey’s multiple comparisons test) (n=3), # indicates p<0.05, significant difference in NKX2.5 expression of iCMs compared to 65-year-old human heart (Student’s t-test) (For clarity, only the statistical comparison of day 21 to all other groups and comparison of NKX2.5 expression to 65-year-old human heart is shown) (C) Immunostaining of F-iCMs at different culture ages for ki67. (Scale bar=20 um) (D) The quantification of ki67 positive cell (%) of F-iCMs and E-iCMs at different culture ages. * represents p<0.05, significant difference when ki67 positive cell (%) on day 21 F-iCMs is compared to F-iCMs at different culture ages. # represents (p<0.05) significant difference when ki67 positive cell (%) on day 21 E-iCMs is compared to E-iCMs at different culture ages. (Student’s t-test) (n≥6)

Another common measure of maturity in iCMs is the low levels of ki67 expression^27^. Although whether cardiomyocytes retain some proliferative capacity or not is controversial^32^, decrease in ki67 levels after birth is a common indicator of exit from the cell cycle, thus of cardiac maturation^33^. Therefore, as another characteristic of maturation, we monitored the change in ki67 expression in iCMs with respect to culture age (Figure 2C, Supplementary Fig. 1A). We observed a gradual decrease in number of ki67 expressing cells with culture age (Figure 2C, D), supporting the previous reports showing a decrease and eventual loss of ki67 expression with age in cardiomyocytes^27,33^. We observed the most drastic decrease starting day 100 of culture where only 3 % of both F-iCM and E-iCM populations were ki67 positive. This percentage did not show a significant difference by day 120 of culture, indicating that ki67 positive cell percentage stabilizes by day 100 of culture. As a control, we examined ki67 expression in human heart sections acquired from 40 and 65-year-old patients. We did not observe any positive nuclear staining for ki67 in correlation with other studies (Supplementary Figure 1B). However, this observation could be due to our small sample size, thus it does not conclusively address the ongoing controversy regarding to what degree the adult heart tissue possesses regenerative capacity.

### Mechanical and Functional Maturation of iCMs with Culture Age

The force generated by cardiomyocytes is one of the most important functional characteristics of the heart tissue. Therefore, we determined the changes in beating velocity of groups of beating iCMs, as well as the beat force of a single iCM within that group, with respect to culture age (Figure 3). The heat maps generated using a custom made MatLab code to quantify the displacement from the brightfield videos of spontaneous beating iCM sheets at different ages of culture demonstrated that beat velocity was highest and most homogenous on day 55 of culture for both cell lines (Figure 3A-D) with a non-significant decrease on day 100.

**Figure 3.**
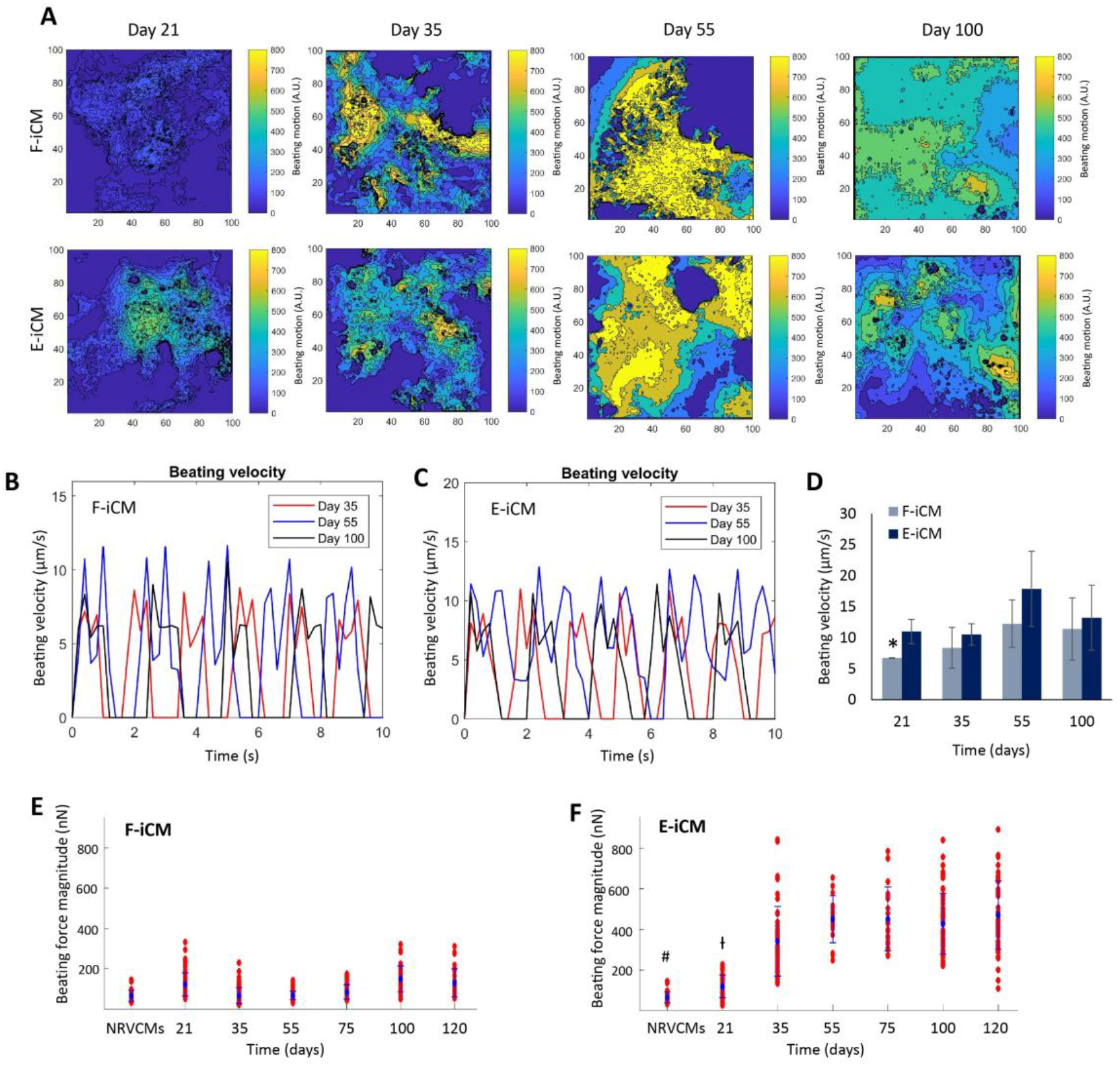
Mechanical characterization of iCMs. (A) The heatmaps showing the beating velocity magnitude (A.U.) and distribution in F-iCMs and E-iCMs at different days of culture. The beat velocity (µm/s) represented with corresponding contraction waveform and frequency on days 35, 55, and 100 of culture (B) of F-iCMs. And (C) of E-iCMs. (D) Average beating velocity (µm/s) of F-iCMs and E-iCMs. * represents p<0.05, statistical significance compared to day 55 of the same cell line (Student’s t-test) (n=5). The beat force magnitude (nN) of (E) F-iCMs and (F) E-iCMs at different culture ages. NRVMs were used as a control. (n=3) # represents p<0.05, beat force magnitude of NRVCMs is significantly lower than iCMs of 35-120 days. Ɨ represents p<0.05, beat force magnitude of day 21 iCMs is significantly lower than iCMs of 35-120 days. (One-way ANOVA followed by Tukey’s multiple comparisons test)

Dwell measurements of single cells within the spontaneously beating iCM sheets using a nanoindenter showed that the highest beat force for F-iCMs was measured on days 21, 100 and 120 of culture ranging from 122.2±56.3-149.6±65.1 nN. We observed that after day 21, E-iCMs were beating stronger compared to F-iCMs (Figure 3B). The highest beat force of E-iCMs was measured from days 55-120 of culture ranging from 427.9±151.0-471.2±168.1 nN. These values are the highest spontaneous beat force values reported for hPSC-derived cardiomyocytes. The highest value reported in other studies showed a total contraction force of hPSCs as 144 nN which did not change over 90 days of culture^34^. This makes the beat force magnitude of E-iCMs the highest reported beat force of hPSC-derived CMs to date, by a 3-fold difference.

To assess the contraction kinetics of the iCMs, we investigated the changes in their Ca^2+^ transient kinetics with culture time (Figure 4A-D) by labeling them with a calcium-sensitive dye, Fluo-4 AM and monitoring the calcium handling during spontaneous beating. The action potential duration (APD) has been shown to prolong with age^6,35^, which presents slower contraction kinetics and slower Ca^2+^ handling, thus the trend in iCMs’ Ca^2+^ transient would tell us how well our in vitro-aged iCM phenotype functionally correlate with adult animal and human heart tissue. Our results showed that as iCMs aged in vitro, the peak to 80% decay was prolonged, consistent with in vivo animal^35^ and human studies^6^ (Figure 4C, D). The fastest contraction kinetics was observed at days 35 and 55 of culture for both F-iCMs and E-iCMs (Figure 4C, D). Both before and after these time points, Ca^2+^ transient decay time was significantly longer. The prolonged APD at 90% repolarization was observed in 14-20-day-old chicken embryos, compared to 1-7-day-old hatched chicks,^36^ suggesting a shortening of APD with maturation. Overall, our results suggest that maturation of iCMs take place through 55 days in culture and the Ca^2+^ handling and contraction kinetics resembling aged human hearts is observed by day 100 in culture.

**Figure 4.**
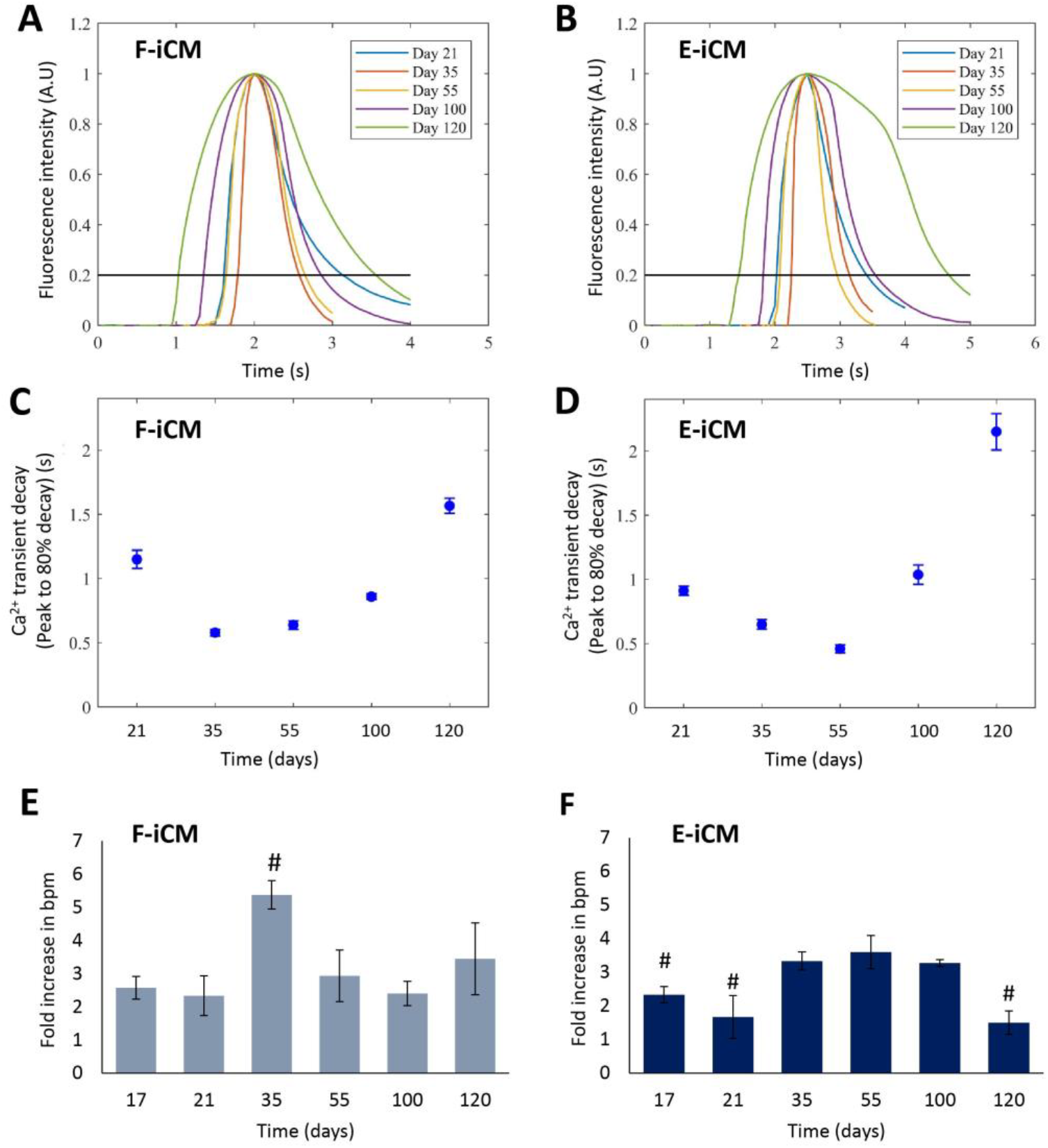
Functional characterization of iCMs. The fluorescence intensity representing the change in Ca^2+^ transient in a single beat in (A) F-iCMs and (B) E-iCMs at different culture ages. The average Ca^2+^ transient decay (peak to 80% decay) in (C) F-iCMs and (D) E-iCMs from day 21 to day 120 of culture. The fold increase in bpm in response to isoproterenol treatment (1 μm) in (E) F-iCMs and in (F) E-iCMs at different days of culture. # indicates p<0.05, significant difference compared to fold increase in bpm on day 55 (One-way ANOVA followed by Tukey’s multiple comparisons test) (n=3).

Finally, we studied the drug response of iCMs from different culture ages using isoproterenol, an epinephrine derivative used in clinic to increase the heart beat rate^37^, that is commonly used for evaluating the functionality of iCMs^26,38,39^. We exposed the iCMs labeled with Fluo-4 AM to 1 µM isoproterenol and recorded the beat rates before and after the exposure (Supplementary videos 4-9). The quantifications showed an increase in beat rate from day 17 to 120 of culture upon exposure to isoproterenol, indicating that the iCMs exhibit physiologically relevant drug responsiveness from early to late stages of in vitro culture, as represented by the increase in spontaneous beat rate (Figure 4E, F). The highest fold increase in spontaneous beat rate was observed on day 35 of culture for F-iCMs (Figure 4E) and at days 35, 55, and 100 for E-iCMs (Figure 4F). Although late stage iCMs were responsive to isoproterenol, both cell lines showed a significantly lower fold increase in spontaneous beat rate by day 120.

Taken together, our characterization results support that iCMs reach a more mature phenotype starting day 35 as shown by the higher beat rate, higher expression of maturity markers, higher beat force and beat velocity, as well as a higher response to isoproterenol, compared to earlier culture ages. Although some of these parameters stabilize after 35 days of culture, a functional deterioration is noticeable by day 75-120 of culture as shown in lowered beat rate, beat velocity, beat frequency, and isoproterenol response. Therefore, we consider iCMs to reach peak maturity between day 35-55 in vitro.

### Characterization of aging phenotype in iCMs with prolonged culture

An important characteristic of aged cells is cellular senescence. We assessed the senescence-associated β-galactosidase (SA-β-gal) activity, as it is one of the most common ways to identify cellular senescence, both in vivo and in vitro^40^. We observed that between day 21-55 of culture, the senescent cell percentage was changing between 1.0 to 5.4% in F-iCMs and E-iCMs (Figure 5A, B, Supplementary Fig. 2). However, on day 100 of culture there was a significant increase in the number of senescent cells, reaching 52.3±3.2% for F-iCMs and 49.2±4.0% for E-iCMs (Figure 5A, B). By day 120 of culture this number increased further reaching 62.5±5.2% for F-iCMs and 57.3±1.0% for E-iCMs. Accumulation of senescent cardiomyocytes with age has been shown in other studies^2,11^. Although to a lesser extent, Chimenti et al. have reported the presence of senescent cardiomyocytes in 76±4-year-old human hearts^11^. They showed that 16% of myocytes and 14% of stem cells found in the aged hearts without any disease condition were senescent.

**Figure 5.**
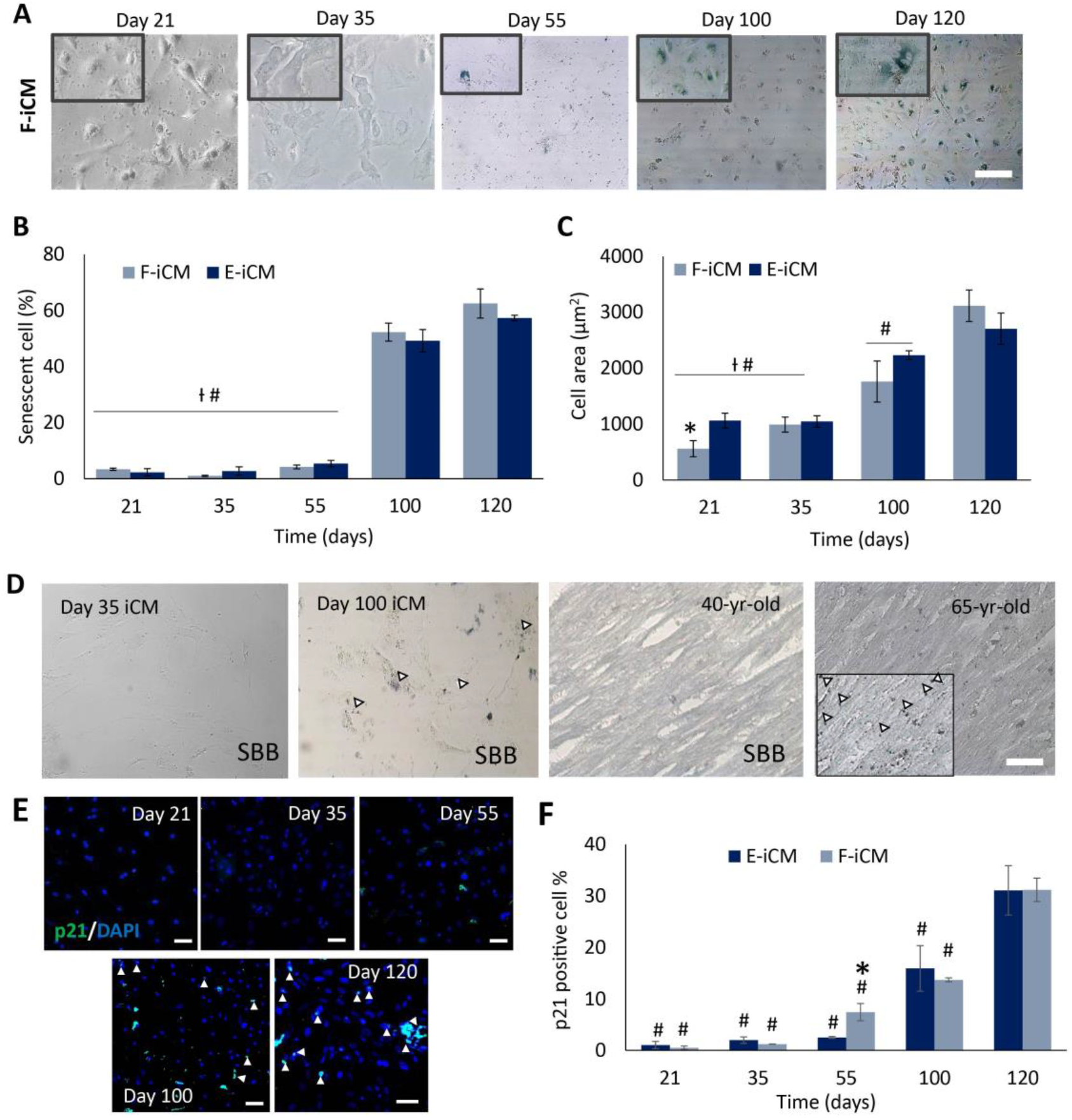
Characterization of aged phenotype of iCMs. (A) The senescence associated β-galactosidase assay images of F-iCMs at different days of culture (blue staining indicates senescence). (B) The senescent cell (%) in F-iCMs and E-iCMs on different days of culture. (Scale bar=200 µm) (C) Cell area (µm^2^) of F-iCMs and E-iCMs at different culture ages. (D) The sudan black b (SBB) staining of F-iCMs on day 35 and 100, and of 40 and 65-year-old human heart sections. (Scale bar=200 µm) (E) Immunostaining of F-iCMs against p21 at different days of culture. (Scale bar=50 µm) (F) Quantification of p21 positive cell (%) of F-iCMs and E-iCMs at different culture ages. * indicates p<0.05, the values for F-iCMs and E-iCMs of same age are significantly different (Student’s t-test), # indicates p<0.05, significant difference compared to day 120 (One-way ANOVA followed by Tukey’s multiple comparisons test), Ɨ indicates p<0.05, significant difference compared to day 120, (One-way ANOVA followed by Tukey’s multiple comparisons test) (n≥6)

Human cardiomyocytes have been shown to enlarge with age in vivo^41^, and the increase in cell area, in general, is another indicator of cellular senescence^42^. We quantified the cell area of iCMs at the early end and later end of 120 day-long culture (Figure 5C). We observed a gradual increase in cell area with culture age. For F-iCMs the cell area increased from 557.9±143.2 µm^2^ to 3116.1±281.7 µm^2^ from day 21 to day 120, respectively.

A similar trend was observed for E-iCMs where the cell area increased from 1061.8±132.2 µm^2^ to 2704±280.4 µm^2^ from day 21 to day 120, showing that the iCMs enlarge with culture time. Considered together with the drop in ki67 expression levels, our results support that iCMs exit cell cycle and enlarge with culture age, resembling the changes human cardiomyocytes go through with age.

Another indicator of cellular aging in the heart is the presence of lipofuscin granules^12,43^. Therefore, we qualitatively determined the presence of lipofuscin granules in iCMs at early and late culture times (Figure 5D), and in human heart left ventricle (LV) sections using a lipofuscin specific stain, sudan black B (SBB)^44^. iCMs were negative for lipofuscin staining at day 35, however, on day 100 of in vitro culture, iCMs had lipofuscin granules as shown by the positive staining for SBB (Figure 5D, left). The human LV sections taken from the 65-year-old heart contained lipofuscin granules (Figure 5D, right). These granules were absent in the human LV sections taken from the 40-years-old heart, indicating that iCMs at day 100 of culture resemble the native aged heart rather than the young heart.

Telomere shortening is another well-known hallmark of aging^45^. Although quiescent, adult cardiomyocytes have shown to go through telomere shortening^5^. An indirect way of observing this phenotype is to monitor p21 expression in cells. The telomere erosion in cells triggers a cellular response that stabilizes tumor suppressor protein p53 and upregulation in p16 and p21 expression. Subsequently, p21 causes cell cycle arrest, causing senescence in proliferating cells^46^. In correlation with this, increase in p21 expression with age has been shown in vivo in higher primates^47^. To further assess the age-related characteristics of our iCMs, we determined the change in p21 expression with respect to culture age (Figure 5E, F, Supplementary Fig. 3). Until day 55 in culture, the p21 expressing cells made up only less than 2% of both F-iCM and E-iCM populations. Although p21 expression significantly increased in F-iCMs starting day 55, reaching 7.4±1.7%, a more drastic increase was observed on day 100 of culture (Figure 5F), reaching 13.7±0.4% for F-iCMs and 15.9±4.4% for E-iCMs. We observed a further increase by day 120 of culture, up to 31.2±2.3% for F-iCMs and to 31.1±4.8% for E-iCMs. Supporting these results, 14% of cardiomyocytes were found to be p16 positive in the aged human heart^11^.

### Fabrication of young and aged myocardial model tissues

Upon thorough characterization of iCMs through prolonged in vitro culture, the cells resembled a young and mature human heart phenotype and functionality between days 35-55 of culture. On the other hand, starting day 100 of culture, the iCMs showed signs of phenotypical and functional deterioration, resembling the aged human heart. Therefore, we defined day 35-55 iCMs as “young iCMs” and day 100-120 iCMs as “aged iCMs.” We then developed 3D, young or aged, cell-laden hydrogel-based myocardial model tissues using the young or aged iCMs, respectively. As the heart ages, not only the cellular content of the tissue changes, but the cellular microenvironment also changes. We measured the stiffness of a 65-year-old heart to be 32±5.9 kPa (Supplementary Fig. 4), which was significantly higher than the stiffness of young hearts (10 kPa)^48^. Therefore, we incorporated the stiffness factor in our model tissues by using 3 different hydrogel compositions with different stiffness: 1) Soft—comprised of methacrylated gelatin (GelMA) (0.4 kPa), mimicking embryonic heart stiffness^49^; 2) Intermediate— arginine-glycine-aspartic acid (RGD) conjugated poly(ethylene glycol) (PEG) (4-arm acrylate, 20 kDa) (PEG-RGD) (8 kPa), mimicking adult heart stiffness^48^; 3) Stiff—PEG (3.5 kDa)-GelMA mixture (P-G) (30 kPa), mimicking aged heart stiffness (Supplementary Fig. 4). It is possible to fabricate tissues with different stiffness using the same hydrogel type, however, that requires changing the fabrication conditions, such as increasing the photoinitiator concentration as well as the UV irradiation dose and time. Since such changes would affect the initial viability of the tissues, we prioritized equalizing the cell viability over hydrogel composition and achieved the abovementioned stiffness values while keeping the fabrication parameters the same. While doing so, we made sure to provide similar levels of cell attachment sequences in all three types of hydrogels. GelMA is gelatin based, thus is rich in RGD sequences. We aimed to have a similar level of cell attachment in the intermediate and stiff hydrogels by incorporating RGD sequences to PEG 4-arm acrylate (20 kDa) and by mixing GelMA with PEG diacrylate (3.5 kDa), respectively. In the case of intermediate tissues, the amount of RGD sequence incorporated (5 µmol/ml) was carefully selected to induce cell attachment at similar levels to collagen gels as shown previously^50,51^. In addition, since achieving a stiffness over 20 kDa with 10% GelMA is not feasible without prolonged exposure to UV (30 minute exposure was shown to yield 36 kPa stiffness)^52^, GelMA-PEG (3.5 kDa) mixture was used. The GelMA concentration in stiff hydrogels was kept the same as used in soft hydrogels, providing the stiff tissues with the same amount of cell attachment sequences. We observed that over 75% of young and aged F-iCMs and E-iCMs, were viable in each hydrogel composition and stiffness used in this study (Figure 6A, B) indicating that the chosen tissue fabrication method is suitable for iCM encapsulation, regardless of iCM culture age.

**Figure 6.**
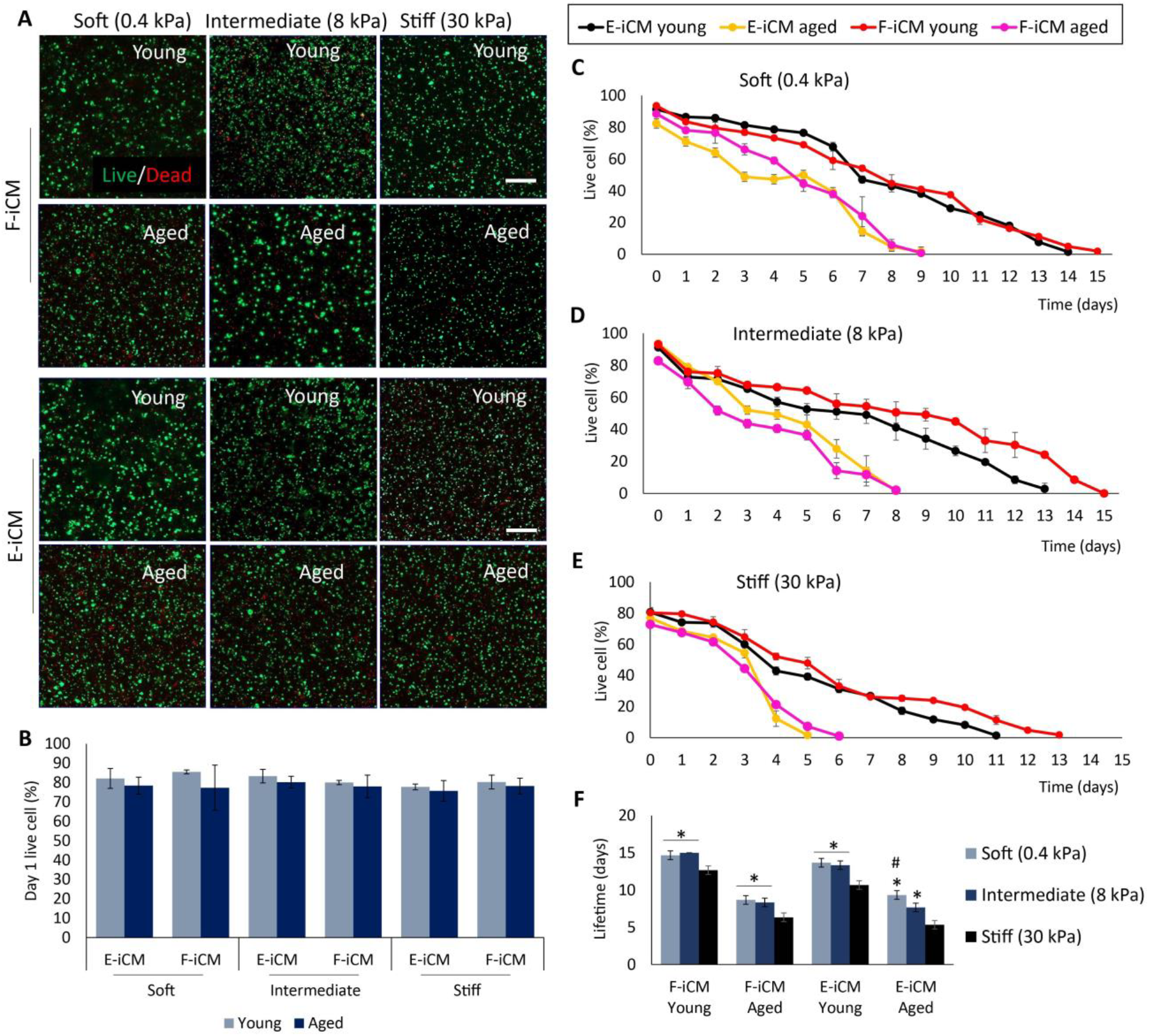
Short term and long-term viability of soft, intermediate, and stiff young and aged tissues. (A) The live/dead images of young and aged tissues 24 hours after encapsulation. (B) The live cell (%) of young and aged tissues on day 1 of culture. The change in live cell (%) with time in (C) soft, (D) intermediate, and (F) stiff young and aged tissues. * indicates p<0.05, significant difference compared to stiff tissue (One-way ANOVA followed by Tukey’s multiple comparisons test), # indicates p<0.05, significant difference compared to intermediate tissue (Student’s t-test) (n≥3) (Scale bars= 200 μm)

### Characterization of the tissue-level aging of the model tissues

We first determined if our tissue models show tissue-level aging characteristics, requisite for developing an in vitro aged tissue model. The fact that the aged iCMs show age-related deterioration on cellular level does not directly translate to aging at the tissue level. One of the most widely accepted definitions of aging at the multicellular level is the exponential increase in the mortality rate by time, referred to as Gompertz Law, which is universal for all organisms complex enough to go through the aging process^53^. In our previous studies^54,55^, we showed that engineered tissue constructs can be made to possess characteristics of organismic level aging, as is associated with a sudden acceleration in mortality rate of cells within the model tissue through the model tissue’s lifetime. Our earlier theory on this tissue level interpretation of the Gompertz law shows that the cellular interdependence within the model tissues and its deterioration by time results in aging behavior as discussed in our recent publications^54,55^. Once the cell loss within one tissue reaches to a critical point, the failure of the remaining cells is accelerated, leading to tissue collapse. Therefore, we investigated whether tissue models incorporated with aged cells show the signs of multicellular aging. To do so, we determined the long-term survival of young and aged model tissues through acquiring the number of dead cells within the tissues, daily. We determined the number of dead cells via ethidium homodimer 1 (EtHD-1) staining every day and calculated the live cell percentage using these values (Figure 6C-F). We observed that in all hydrogel compositions the viability of young tissues was higher compared to aged tissues. We also observed a significant effect of stiffness on long-term tissue viability. The lifetime of stiff tissues was lower by 2-3 days compared to soft and native stiffness tissues. This suggests that not only the cell age but also the microenvironment stiffness is crucial for tissue aging. Importantly, we observed a collapse in the survival curve of only the aged, stiff tissues. This specific pattern of change in tissue viability by time is the abovementioned characteristic failure mode observed only in complex, aging systems^54^. This observation crucially shows that using in vitro aged cells in combination with a matrix of the age-appropriate stiffness we can develop physiologically relevant aged model tissues suitable for studying age-related diseases.

### Stress response of young and aged myocardial tissue models

The functional and phenotypical deterioration in aged iCMs make them a good candidate to be used in building aged tissue models. However, the impairment in cell function might not be enough to correctly mimic the aged tissue/organ phenotype. As such, we first determined the effect of in vitro tissue age and stiffness on survival under MI mimicking conditions, specifically in response to reperfusion injury (RI). The immediate action taken after MI is to reoxygenate the infarct tissue by restoring blood flow. However, the sudden flow of oxygen-rich blood to the ischemic area induces oxidative stress and often results in further cell death causing RI^56–58^. We tested 3 different RI conditions: 1) 16 h H_2_O_2_ exposure followed by 2 h normoxia to investigate tissue response to oxidative stress followed by normoxia, exemplifying a commonly used in vitro RI model (Figure 7A, B, Supplementary Fig. 5); 2) 48 h hypoxia (1% O_2_) followed by 24 h normoxia (21% O_2_) to investigate the late tissue response to reoxygenation (Figure 7C, D, Supplementary Fig. 6); 3) 48 h hypoxia (1% O_2_) followed by 2 h normoxia (21% O_2_) to investigate the early tissue response to reoxygenation (Figure 7E, F, Supplementary Fig. 7). Subsequently, tissue survival was calculated using Live/Dead assay. The tissue viability after stress exposure was normalized to respective normoxia controls, and the results were represented as normalized survival. We observed that the survival of aged tissues was significantly lower than that of young tissues regardless of the cell type or stiffness when exposed to 48 h hypoxia+24 h normoxia or 16 h H_2_O_2_+2 h normoxia. E-iCM tissues that were exposed to 16 h H_2_O_2_ followed by 2 h normoxia showed a lower survival compared to E-iCM tissues that were exposed to 48 h hypoxia followed by 24 h normoxia.

**Figure 7.**
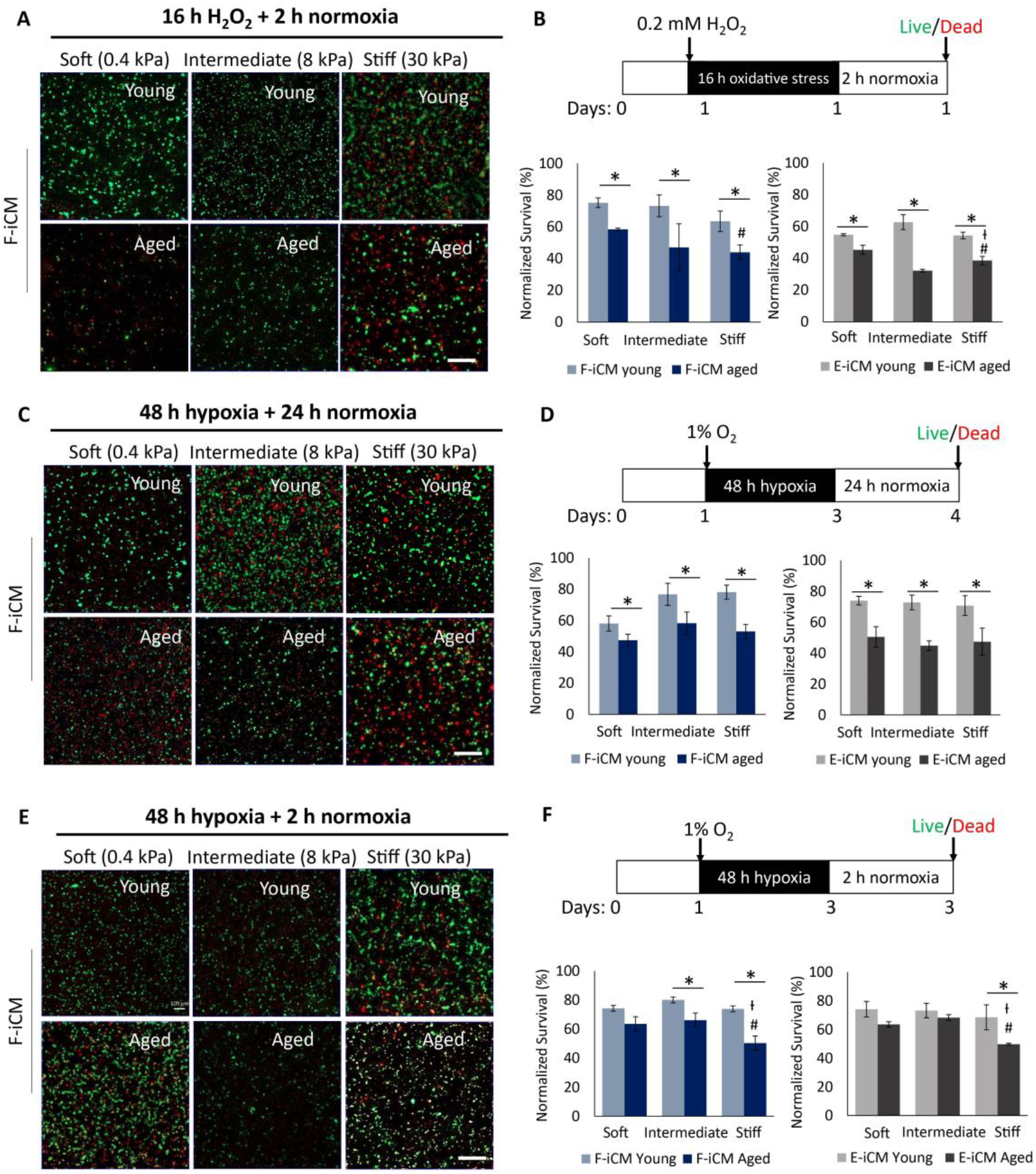
Stress response of young and aged, soft, intermediate and stiff tissues. (A, C, E) Live/dead images and (B, D, F) normalized survival (%) of young and aged tissues with different stiffness after exposure to (A, B) 16 h H_2_O_2_ + 2 h normoxia, (C, D) 48 h hypoxia + 24 h normoxia, (E, F) 48 h hypoxia + 2 h normoxia. * indicates p<0.05, significant difference compared to young tissue at same stiffness, student’s t-test, # indicates p<0.05, stiff tissue survival is significantly different than soft tissues (One-way ANOVA followed by Tukey’s multiple comparisons test), Ɨ indicates p<0.05, stiff tissue survival is significantly different than intermediate tissues (One-way ANOVA followed by Tukey’s multiple comparison’s test) (n≥3) (Scale bars=200 μm)

Interestingly, under 48 h hypoxia+2 h normoxia conditions, the most significant age-related difference in survival was observed with stiff tissues, suggesting that stiffness is crucial in stress response regulation. In addition, there was a non-significant difference in survival of young vs. aged E-iCM tissues at soft and intermediate stiffness after 2 h normoxia that followed 48 h hypoxia (Figure 7F). We also observed a difference in response to long (24 h) vs. short (2 h) term normoxia following 48 h hypoxia. While the survival of aged tissues exposed to 24 h normoxia was significantly lower than young tissues exposed to the same stress, this was not the case for tissues exposed to 2 h normoxia. This potentially shows that short periods of normoxia subsequent to hypoxia can be tolerated in in vitro aged tissue models or that 2 h normoxia is not long enough to affect the cell viability significantly. However, further studies are required to gain a detailed understanding of the effect of short term vs. long term reperfusion injury in in vitro aged tissue models.

### Tissue level functional deterioration is observed in aged tissues

Although the age factor is mostly neglected, there are in vivo and in vitro studies showing that at later ages the otherwise beneficial effect of therapeutics would be impaired. Inhibiting the soluble epoxide hydrolases whether using inhibitors (EHI) or genetic manipulations, has been shown to hold promise to decrease infarct size and induce cardioprotection in vivo^59–62^. A recent study by Jameison and colleagues, however, showed that the cardioprotective effects of genetically deleting epoxide hydrolases was significantly impaired in aged mice, compared to young animals. Therefore, an in vitro aged tissue model should not only possess cell and tissue level aging phenotype and related characteristics but should also mimic the pharmacological differences observed in vivo. We, therefore, investigated the effect of soluble epoxide inhibition on survival and ROS production in our young and aged tissue models under RI mimicking stress. To achieve that, we supplemented the culture media with EHI during hypoxia and the subsequent normoxia treatment. In order to capture the ROS levels, we have selected the 48 h hypoxia followed by 2 h normoxia stress treatment as the ROS levels were undetectable after 24 h of normoxia, due to short lifetime of oxygen radicals. We observed that EHI presence has improved the survival of young tissues regardless of microenvironment stiffness (Figure 8A-C, Supplementary Fig. 8). In addition, at soft and intermediate stiffness, young tissues supplemented with EHI had significantly lower ROS levels compared to the young tissues without the supplement (Figure 9A, B, Supplementary Fig. 9). Statistically significant improvement in survival and the reduction in ROS levels, however, were not observed in soft and intermediate stiffness aged tissues, indicating an age-related impairment in stress response on tissue level. In stiff tissues, we observed a similar trend in both survival and ROS production. Although at this stiffness the cardioprotective effect of EHI yielded a significant increase in aged tissue survival (Figure 8C), the improvement in survival of young tissues was more noticeable. The 50.3±2.1% (F-iCM) and 49.8±4.9% (E-iCM) cell survival in aged tissues increased to 65.4±2.2% (F-iCM), and 64.2±2.9% (E-iCM) with EHI supplementation. On the other hand, the 73.9±4.5% (F-iCM) and 68.5±8.1% (E-iCM) cell survival in young tissues increased to 98.2±4.9% (F-iCM) and 94.6±3.2% (E-iCM) showing that in vitro aged tissue models reflect the age-related reduction in drug sensitivity observed in human hearts (Figure 8C). Interestingly, the ROS production of young and aged tissues was comparable at aged heart stiffness without EHI. Young tissues had significantly lower levels of ROS only when supplemented with EHI (Figure 9C). Although the survival trend was similar among the 3 different stiffness values and the 2 different cell lines (Figure 8A-C), ROS production in young tissues was similar to that of aged tissues only in stiff tissues.

**Figure 8.**
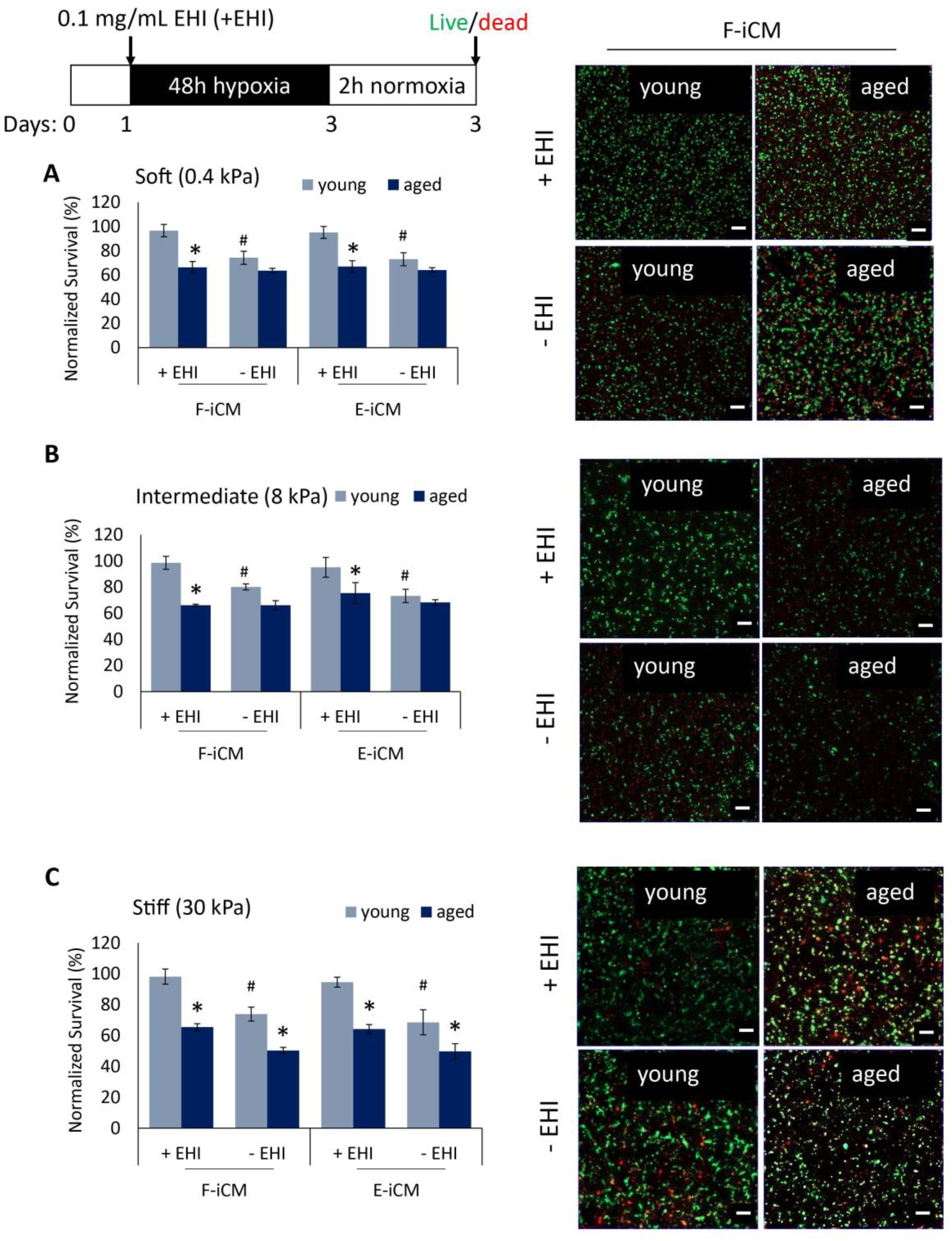
Effect of EHI on survival of young and aged tissues at different stiffness under 48 h hypoxia + 2 h normoxia. The normalized survival (left) and corresponding live/dead images (right) of (A) soft, (B) intermediate, and (C) stiff young and aged tissues with and without EHI (+EHI and –EHI, respectively). * indicates p<0.05, aged tissue survival is significantly different than young tissue under same condition (Student’s t-test), # indicates p<0.05, young tissue survival is significantly different under -EHI vs +EHI conditions (Student’s t-test) (Scale bars= 100 μm)

**Figure 9.**
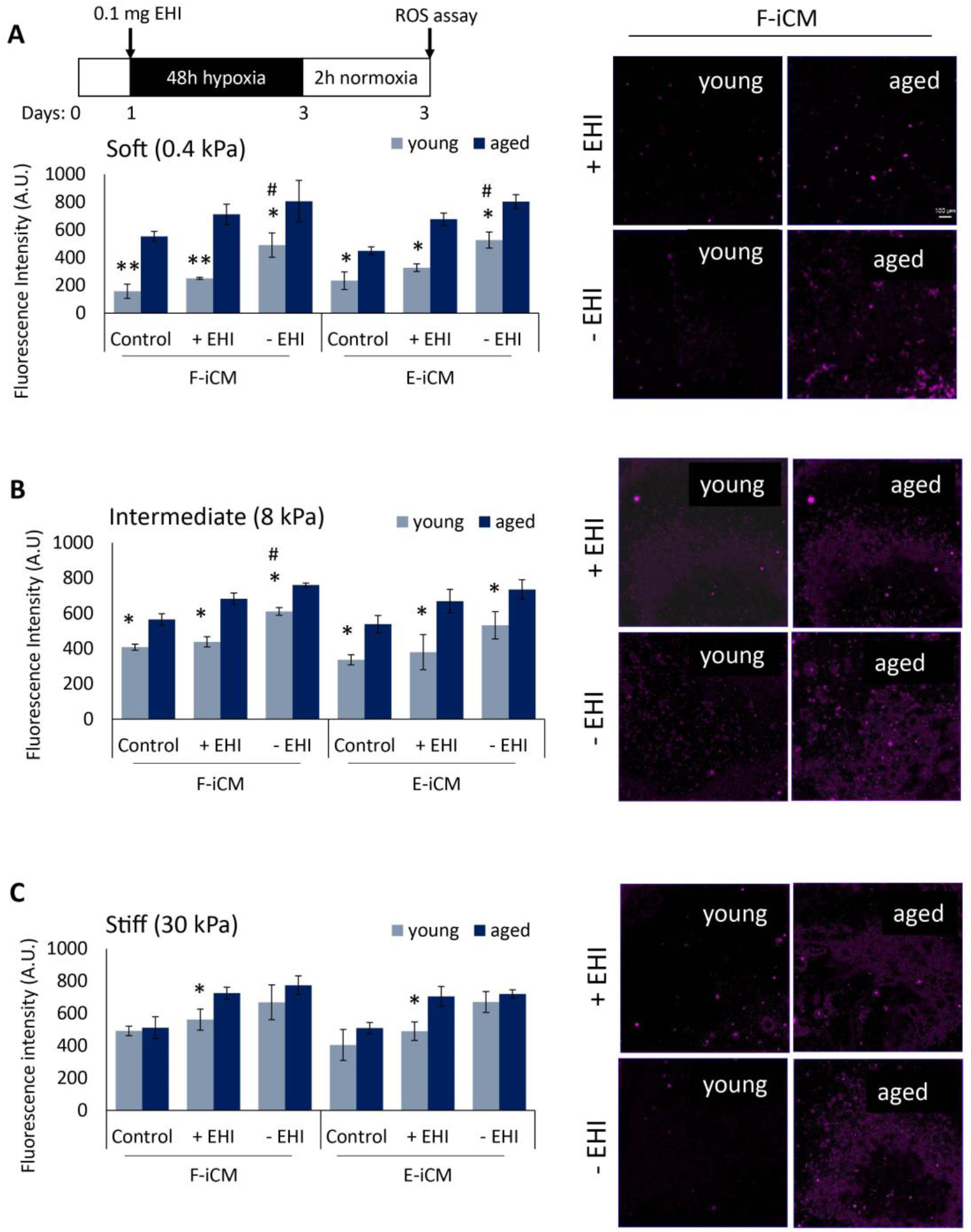
Effect of EHI on ROS production in young and aged tissues at different stiffness under 48 h hypoxia + 2 h normoxia. The ROS level quantified as the fluorescence intensity measurements (left) and corresponding ROS assay images (right) of (A) soft, (B) intermediate, and (C) stiff young and aged tissues with and without EHI (+EHI and –EHI, respectively). * indicates p<0.05, ** indicates p<0.01, ROS levels is significantly different than aged tissues under the same conditions (Student’s t-test), # indicates p<0.05, ROS levels in young tissue is significantly different under -EHI vs +EHI conditions (Student’s t-test) (Scale bars= 100 μm)

Overall, we determined that our aged tissues have a lower survival and higher ROS production upon exposure to RI mimicking stress conditions, compared to young tissues. Importantly, the difference in survival between young and aged tissues was higher at aged heart stiffness, suggesting that not just the age of the cells but also the mechanical characteristics of cells’ microenvironment must be considered to develop an age-appropriate tissue/disease model, as we have shown here.

## Discussion

Aging is a well-known contributor of CVD, however, majority of studies investigating CVD are done using young animals^19^ or cells although the physiological and pathological differences of young and aged tissues/cells are well-documented^2,3,6,8,63^. Use of age-appropriate animals can be costly and time consuming and the physiological differences between animal and human pathology still presents a challenge. Alternatively, using tissue engineering techniques and hiPSCs it is now possible to develop human-origin tissue models with controlled microenvironments. Although many examples of incorporating hiPSC-derived cells to engineered tissues exist^21,38,64,65^, the aged phenotype of the myocardial cells or the tissue has not been considered to date. In this study, we developed the first in vitro aged cardiac tissue model with aged myocardium stiffness-mimicking ECM incorporated with age-appropriate hiPSC-derived cardiomyocytes. This model provides a highly controlled platform for in vitro cardiac tissue modeling that is: (1) human-origin, (2) time and cost effective, and (3) pathologically relevant (both biologically and mechanically). Moreover, this model can potentially be modified to develop in vitro aged disease models to study other age-related diseases.

While hiPSC-derived cardiomyocytes hold promise for human-origin and patient specific disease modeling, it has been widely reported that these cells resemble the fetal cardiomyocytes rather than adult ones^33^. It has been shown in other studies that iCMs mature with time^66^ and with mechanical^33,67^ or electrical^33,67,68^ stimuli. Here we show that regardless of the iPSC origin, iCMs mature in vitro with prolonged culture time, and start showing marks of cellular aging at the later culture stages, through 120 days of culture we conducted in this study. This is consistent with other assessments which report maturation of iCMs cultured up to 120 and 180 days^66,69^. Although iCMs start beating by day 10 and show striated TNNT2 structures as early as day 17, starting day 35 of culture they showed superior beat rate. The increased maturity of iCMs with culture age was also reflected in their expression profile with a consistent increase in mRNA levels of maturity markers such as CASQ2, MYH6, MYL2, S100A1, and TNNT2. In support of this, the expression of markers such as ki67 and NKX2.5, which are known to decrease with age in vivo^31,70^ and in vitro^27,33^, showed a gradual decrease with culture time. Unlike results reported by Lundy et al.^69^, iCMs used in this study reached peak molecular maturity by day 55 of culture, compared to day 80, and beyond this culture age we observed a leveling off in maturity marker expression. The slower contraction mechanics they observed at later stages, however, agree with our findings and suggests the start of functional deterioration with prolonged culture time.

In addition to molecular and phenotypical characteristics, the contraction mechanics of iCMs showed an increase with culture time, reaching a peak between day 35-55, followed by a decrease with longer culture as shown by the prolonged Ca^2+^ transient decay after day 100. Age-related impairment of contraction mechanics and electrophysiological function in cardiomyocytes has been reported in vivo in animals^6,63^ and humans^7,71^ as presented by a lengthening of APD and slower Ca^2+^ transients. In other studies, this physiologically relevant age-driven prolongation in APD of iCMs has not been reported. In one study, the APD at 90% repolarization values for late-stage hiPSC and human embryonic stem cell-derived cardiomyocyte cultures has shown to remain constant through 120 days of culture^69^ and APD has not been determined in other long-term culture studies^66^. Similarly, the peak to 90% decay in Ca^2+^ has been shown to be improved after day 80 in culture^66^. This difference can be attributed to the later maturation (day 80-120 of culture) observed in the mentioned study, potentially delaying the effects of aged cell phenotype. Similarly, a thorough mechanical characterization of iCMs with prolonged culture times has not been conducted before. Here, we show that the beat velocity of iCMs reach its peak at day 55 and was higher than that of other iCMs reported,^38,39^ and showed a non-significant decrease by day 100 in culture consistent within the two different hiPSC lines used. Importantly, we observed a beat force magnitude of over 450 nN, which is the highest beat force magnitude for hiPSC-derived cardiomyocytes reported in the literature. Although fibroblast origin iCMs showed a significantly lower beat force, it is still comparable to or greater than that of neonatal rat cardiomyocytes (NRVCMs) and among the highest reported in the literature^72^. In addition, we show that F-iCMs are responsive to isoproterenol and had previously shown that they are electrically responsive and can be paced using micro electrode array systems^21^. The difference between the beat force of F-iCMs and E-iCMs can be attributed to their origin. It is documented that iPSCs retain epigenetic memory to some extent^73,74^. Therefore, the endothelial origin E-iCMs potentially retained the epigenetic signature of endothelium, which in turn aided yielding iCMs with superior beat force since both endothelium and cardiomyocyte lineage are derived from mesoderm. However, this claim needs to be investigated in detail with further studies.

We also investigated the age-related changes in responsiveness to β-adrenergic receptor stimulation. We showed the age-related reduced drug response in iCMs through their impaired response to isoproterenol. In one study, White and Leenen reported that response to adrenaline was significantly reduced in individuals of age 60±2, compared to individuals of age 30±2^14^. A similar difference of drug response in young vs. aged individuals was reported in other studies as well^9,10^.

Overall, the consistent trend of improvement in maturity with culture time followed by a functional deterioration at later stages of culture led us to investigate the late stage iCMs further and compare them to aged human tissue. We observed that iCMs show basic characteristics of cells comprising aged tissues. There is still no single phenotype that defines an aged cell. Aging is commonly defined as time-dependent functional deterioration^45^. At the cellular level, senescence, telomere shortening, DNA damage, and ROS accumulation are listed as some of the hallmarks of aging ^47,75^. In agreement with the functional deterioration we observed with decrease in beat velocity and frequency, and prolonged Ca^2+^ transient decay, the iCMs showed higher levels of senescence and gradual increase in p21 expression with culture time. In addition, increase in cardiomyocyte cell area has been correlated with aging^70^, thus supporting our claim that iCMs resemble native aged human cardiomyocytes. Importantly, presence of lipofuscin granules, which are composed of lipid-containing residues as a result of lysosomal digestion, is considered an aging marker and has been reported to be present in aged human hearts^12,43,44^. We report here for the first time that long term culture of iCMs lead to the presence of lipofuscin granules and this phenotype distinctly resembles the 65-year-old human heart, as we did not detect these granules in 40-year-old human heart.

iPSC-based cardiovascular disease models^64^, both 2D^65,76^ and 3D^21,27,77^, have been designed for various applications. It is extremely crucial to replicate the disease pathology as closely as possible. Therefore, studying age-related diseases such as RI require the use of age-appropriate animal or in vitro models. Accelerated aging of proliferative cells in vitro have been shown by our group^54^ and others^78,79^ previously. In the case of iCMs, however, aging, defined with senescence in most studies, is not induced due to proliferative exhaustion or oxidative stress. As it is well-established that cardiomyocytes do not proliferate extensively, and as supported by the decrease in ki67 expression with age, late stage iCMs used in this study do not simply reach senescence due to reaching their proliferative limit. Therefore, as supported by the age-related characterization we show here, we strongly believe that iCMs past day 100 of culture are possible candidates for developing human-origin, age-appropriate disease models to study age-related diseases. In addition to providing the advantage of using human cells, our in vitro aged tissue models are time effective models. Where an age-appropriate mouse model would require over 20 months of aging, we can achieve aged iCMs within 4 months. We obtain cells that resemble aged human cells approximately 200 times faster than the in vivo aging process. This accelerated aging in vitro can be attributed to many factors such as the lack of supporting cells, lack of stem cells, and lack of the systemic regulation that are present in vivo, as well as high oxidative content of in vitro culture conditions which would create a baseline oxidative stress that is not seen in native tissues until late ages.

Although achieving the correct cell phenotype is of upmost importance, cell phenotype is not the only factor that contributes to organ/tissue function. The microenvironment of heart also shows changes with age. Several reports show that both the animal hearts^3^ and human^80,81^ hearts stiffen with age. Therefore, we developed a tissue model where we incorporate the iCMs that show aged cell phenotype in matrices with stiffness mimicking that of aged human hearts. We also developed tissues that are very soft, resembling embryonic heart stiffness, and ones resembling young heart stiffness and investigated the effect of tissue stiffness on tissue function and survival. Although cells were highly viable initially, regardless of their culture age and tissue stiffness, increased stiffness negatively affected the lifetime of the tissues resulting in cell death in stiff tissues sooner than soft tissues. More importantly, when we examined the functional form of the lifetime measurement curves (i.e. the number of viable cells each day of the culture until all the cells are dead) our results showed that the viability of the model tissues with aged iCMs and high tissue stiffness resulted in lifetime curves that are characteristic of aging systems, as indicated with a sudden collapse in the cell viability during the long-term tissue culture corresponding to increase in probability of death over time. In model tissues composed of young iCM with all tested stiffness values and in softer tissues with aged iCMs, the decay of the tissues was more gradual, and the aging-characteristic collapse was not as prominent.

An important age-related alteration in cardiovascular physiology is the impaired stress response mechanism of cardiomyocytes and the related reduced effect of therapeutics^10,13,14,82^. Therefore, an aged disease model should reflect these important changes. The consistent lower survival of our aged tissues with exposure to 16 h H_2_O_2_+2 h normoxia and 48 h hypoxia+24 h normoxia, regardless of stiffness, suggest that using aged iCMs provides a better in vitro model compared to using young iCMs. This is consistent with other in vivo^3,8,63,75^ reports showing that there are significant differences between young and aged heart physiology and drug response, leading to reduced cardioprotection under stress conditions. Importantly, under 48 h hypoxia + 2 h normoxia treatment, the aged tissues with soft and intermediate stiffness coped with stress almost as well as the young tissues. On the other hand, we observed significantly lower survival rates in aged tissues, compared to young tissues in stiff hydrogels. This shows, in addition to the cell age, the effect of tissue microenvironment is crucial. This supports our findings showing the unique aged tissue failure behavior of tissues with the pathologically correct cell age and stiffness.

We further studied the resemblance of our in vitro aged tissues to aged animal and human heart. To do so, we treated our tissues with EHI, the cardioprotective effects of which towards myocardial IR injury have been well documented^59,61,62^. EHIs contribute to cardioprotection by avoiding epoxy eicosatrienoic acids (EETs) from getting further metabolized. EETs are derived from arachidonic acid through metabolism and they were shown to have anti-inflammatory, anti-hypertensive and anti-apoptotic effects^83^. In animal studies perfusion with EETs improved recovery following RI and reduced infarct size^83–85^. Consistent with these results, genetic deletion of epoxide hydrolases or use of EHIs have been shown effective in improving LV pressure, decreasing infarct size and fibrosis, and reducing cardiomyocyte death^59–61,86,87^ However, recently Jameison et al., has shown that the protective effect of genetic deletion of soluble epoxide hydrolase was impaired in 16-month-mice, compared to 3-month-old ones upon myocardial infarction^88^. Similar to in vivo results, effect of EHI was impaired in our stiff human-origin in vitro aged tissue models upon RI mimicking stress. Most strikingly, with soft and intermediate tissues we did not observe any improvement in the survival of aged tissues. The impaired yet significant improvement in aged tissue survival was present only at stiffness mimicking aged hearts, further supporting the conclusion that creating the age-appropriate cell phenotype and microenvironment is crucial to developing a successful disease model.

Although crucial, determining the decrease in tissue survival is not the only measure of aging. Mitochondrial dysfunction is another hallmark of aging and is highly correlated with CVD pathology^89,90^. Healthy mitochondrial function is essential in heart function and survival and is required to withstand RI^89–92^. In aged hearts, however, mitochondria has been shown to have disrupted morphology and has increased production of ROS^90,91^. Owing to their reactive nature, ROS oxidize various biomolecules and inactivate enzymes reducing the functional ability of the heart. Therefore, aging contributes to the increased susceptibility to CVD, and diminishes the effectiveness of cardioprotective strategies. Our in vitro aged tissues consistently had higher ROS levels, regardless of the cell type and tissue stiffness. In addition, the impaired cardioprotective effect of EHI in aged tissues was reflected on ROS production as well. The aged tissues treated with EHI had lower levels of ROS, compared to tissues that did not receive the treatment. However, this decrease in ROS levels was non-significant. On the other hand, the young tissues showed a significant decrease in ROS levels with EHI treatment, in line with the observations reported in vivo. Interestingly, young stiff tissues showed higher ROS levels compared to soft and intermediate young tissues. NRVCMs have been shown to form unaligned sarcomeres and stress fibers at stiffness values higher than native heart stiffness^93^. Similarly, young iCMs potentially experienced more stress compared to soft and intermediate tissues, causing elevated ROS levels. This correlates well with the decreased lifetime of young stiff tissues compared to lifetime of soft and intermediate tissues.

Overall, here we showed a thorough characterization of human-origin iCMs and reported the highest beat force yet achieved. In addition, we showed for the first time that although iCMs gain a more mature phenotype with prolonged culture age, after about 100 days they start to experience functional deterioration, much resembling the aged human heart cells. Importantly, the aging observed in cellular level is translatable to tissue level by incorporating these aged iCMs to biocompatible matrices with age-appropriate stiffness. Only at stiffness values resembling that of an aged heart, the tissue models of aged cells behave similar to the hearts of aged animals and humans. This provides a novel in vitro model of aged myocardial tissue that is: 1) time efficient, because aging process takes only 4 months, 2) physiologically relevant, because it is comprised of human cells, and 3) pathologically relevant, because aging characteristics are observed both at the cell and tissue level. We believe that although the absence of supporting cells and the overall systemic support causes iCMs to age much faster in vitro, these cells and the respective engineered tissue models present a valuable platform for intermediate studies where scientists can validate the findings from animal studies, before investing vast amounts of money and time in the follow up clinical studies.

## Materials and Methods

De-identified human heart samples were acquired from Indiana Donor Network under an ongoing IRB approval (Prof. Keith March, University of Florida) for deceased donor tissue recovery. Two hearts, ages 40 and 65, that were disqualified for transplantation were collected and cut into small pieces. The information on the patients is presented in Supplementary Table 1.

### Human heart sample fixation and sectioning

Human heart samples were fixed with paraformaldehyde and 10 μm thick sections were taken using a cryomicrotome (Thermo Fisher). The sections were immediately transferred to superfrost glass slides and maintained at −80°C until used.

### Cardiomyocyte differentiation

We used 2 different hiPSC lines from two different sources: DiPS 1016SevA (from skin fibroblasts)^24^ and Huv-iPS4F1 (from human umbilical cord vein endothelial cells (HUVECs))^25^, for cardiomyocyte differentiation. The hiPSCs were cultured on geltrex (Life Technologies) coated tissue culture flasks in mTeSR-1 medium (StemCell Technologies). At approximately 80% confluency, the hiPSCs were collected using Accutase (StemCell Technologies) and seeded on geltrex coated glass coverslips using mTeSR-1 supplemented with 5 µM Rock inhibitor (StemCell Technologies) at 1.5×10^5^ cells/cm^2^. Differentiation to cardiomyocytes was achieved using a previously established protocol^26^. Briefly, on day 1, the hiPSCs medium was replaced with RPMI 1640 (Life Technologies) medium supplemented with B27 supplement without insulin (B27(-)) (Invitrogen) and with 10 µM GSK-3β inhibitor, CHIR (Stemgent). After 24 hours this media was replaced with RPMI 1640 supplemented with B27(-). 48 hours after the medium was replaced with RPMI 1640 supplemented with B27(-) with the addition of wnt inhibitor IWP4 (Stemgent). On day 6 the media was replaced with RPMI 1640 supplemented with B27(-). Starting day 9 the cells received RPMI 1640 supplemented with B27 supplement with insulin (B27(+)) (Invitrogen). From this point on the media was changed every 2-3 days with fresh RPMI 1640 supplemented with B27(+). Beating cardiomyocytes were observed starting day 10 of differentiation.

### Structural characterization and immunofluorescence

The beating efficiency per differentiation and the spontaneous beat rate of iCMs were determined by visual observation and through video recordings at different days of culture, respectively. FACS against TNNT2 was used to quantitatively determine the percentage of iCMs in a differentiation population. FACS analysis was done following a previously established protocol^94^. Rat cardiomyocytes were used as positive controls and for gating, and unstained cells were used as negative control and for gating.

Immunostaining against TNNT2 was performed to qualitatively measure the differentiation efficiency, as well as to confirm the striated troponin structure, an indicator of maturity in iCMs. Ki67 immunostaining was performed to determine percentage of ki67 positive cells over culture time. Briefly, the iCMs were fixed at different days of culture with paraformaldehyde (4%, 10 min, RT) followed by permeabilization with triton-X 100 (Sigma Aldrich) (0.1%, 15 min, RT) and blocking with 10% goat serum (Invitrogen) (1 h, RT). The samples were then incubated with the respective primary antibodies for TNNT2 (Abcam, ab45932) and ki67 (ThermoFisher, 14-5698-82) (overnight, 4°C) and then with the species-appropriate secondary antibodies (Life Technologies) (6 h, 4°C) with a thorough wash in between. The respective images were acquired using a fluorescence microscope (Zeiss, Hamamatsu ORCA flash 4.0).

### Quantitative Real Time PCR (qRT-PCR)

RNA was collected from iCMs at different days of differentiation using a total RNA isolation kit (RNeasy, Qiagen). Fixed human heart tissue samples were first snap-frozen in liquid nitrogen, homogenized using mortar-pestle and then lysed in Trizol. After tissue debris was separated through centrifugation, the supernatant was mixed with chloroform and shaken vigorously for 30 sec. The solution was then centrifuged following the incubation step for phase separation. RNA was collected from the upper layer formed after centrifugation, using a total RNA isolation kit (RNeasy, Qiagen). RNA purity and concentration were measured on a NanoDrop 2000 Spectrophotometer (Thermo Fisher Scientific). RNA was converted into cDNA using the iScript cDNA Synthesis Kit (Bio – Rad). Validated human gene-specific primers were purchased from Bio-Rad (Supplementary Table 2). iTaq SYBR Green Supermix (Bio-Rad) was used for qRT-PCR reactions and run on CFX Connect 96 Real-Time PCR system (BioRad) in triplicates. The relative expression of genes was quantified by the ΔΔCt method using GAPDH as the housekeeping gene.

### Video analysis of iCM contractility

Block-matching algorithm^95^ was performed using MATLAB to analyze contractility of iCMs in this study. Briefly, the bright field videos were exported as a series of single-frame images files. The images were then divided into square blocks of N×N pixels. Movement of a given block at the k^th^ frame is tracked by matching its intensity to an identically-sized block of pixels in the (k + 1)^th^ frame within a square window of width N + 2w. The values of n and w were determined as 16 and 7, respectively, in this study based on the speed of calculation and accuracy of the block-matching method. The matching criterion used for block movement is Mean Absolute Difference (MAD) as given below:

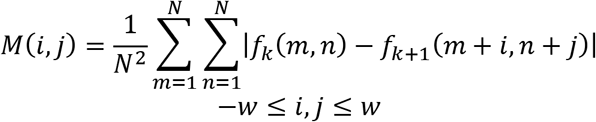

where *f*_*k*_(*m*, *n*) is the intensity at coordinates (*m*, *n*) of a given block at the k^th^ frame and *f*_*k*+1_(*m* + *i*, *n* + *j*) represent the intensity at new coordinates (*m* + *i*, *n* + *j*) of the corresponding block at the (k + 1)^th^ frame. The movement of the block was determined as *min*(*M*(*i*, *j*)).

### Contractile force measurements

The contractile forces of iCMs and 65-year-old human heart tissue were measured with the nanoindenter through dwell experiments as shown previously^4^. Briefly, the nanoindenter probe was brought into contact with the samples, and the probe’s displacement was kept constant (i.e. probe was dwelled on the sample) for 30 seconds to dynamically measure its deflection, which is proportional to the cell’s contractile force along the transverse direction with respect to the substrates. The probe used had a spring constant and tip diameter of 0.068 N/m, and 19 µm, respectively. A customized MATLAB code was developed to separate each single beat and to calculate averaged contractile forces.

### Ca^2+^ transient measurement

A customized MATLAB code was developed in this study to measure Ca^2+^ transient in iCMs at each timepoint. First, maximum value of each intensity peak was determined from intensity vs time curves acquired from videos of iCMs labeled with the calcium sensitive dye, Fluo 4, at different times of culture. The decay in Ca^2+^ transient was then measured as the duration for the intensity to reduce to 80% of its maximum (peak to 80% decay). Next, the intensity was normalized to its maximum value and the overlapped projection of the beats from each video was plotted.

### Determining isoproterenol response of iCMs

iCMs at different days of culture were labeled with Fluo-4, a calcium-sensitive dye (Life Technologies), as instructed by the manufacturer. Videos were recorded using a fluorescence microscope (Zeiss, Hamamatsu ORCA flash 4.0) right before incubation with isoproterenol to determine the baseline for calcium transient and beat rate of the cells. Then medium of the cells was replaced with medium supplemented with 1 µM isoproterenol and incubated for 10 min at 37°C. Immediately after isoproterenol incubation the beat rate and calcium transient of iCMs were recorded using a fluorescence microscope (Zeiss, Hamamatsu ORCA flash 4.0).

### Determining senescence in iCMs

Senescence in iCMs was determined using SA-β-gal assay and sudan black B staining. SA-β-gal assay was performed following manufacturer’s instructions (Millipore). Sudan black B staining was performed to determine lipofuscin granules in the cells and human heart tissues. The iCMs at were fixed with 4% paraformaldehyde at day 35 (young) and day 100 (aged) of culture. Sudan black B was dissolved in 70% ethanol with overnight vortexing to make up a 1% (w/v) solution. This solution was then diluted with PBS to a 0.05% solution, which was applied to the cells (5 min, at RT). The cells were then washed with PBS three times and imaged using a bright field microscope (Leica). Fixed human heart tissue sections were first incubated with PBS to hydrate the tissue (5 min, at RT). Then the tissue sections were incubated with 0.05% sudan black B solution for 5 min at RT. After staining the tissue sections were washed extensively with PBS and images using a bright field microscope (Leica).

Immunostaining against p21 was performed following the protocol described in “Structural characterization and immunofluorescence” section using a primary antibody specific for p21 (Abcam, ab54562) and a species-appropriate secondary antibody.

### Determining cell area

To measure the cell area, six images of iCMs at different days of culture were used. The circumference of at least 20 cells in each image were selected using imageJ and the area (µm^2^) was determined by setting the imageJ scale to the known scale in the images (µm). The results were represented as average ± standard deviation.

### Young and aged tissue fabrication

Young (day 35-55) or aged (day 100-120) iCMs were collected using trypsin-EDTA and encapsulated in 3 different hydrogels; 1) methacrylated gelatin (GelMA), 2) PEG (4-arm acrylate, 20 kDa, JenKem) modified with cell attachment peptide arginine-glycine-aspartic acid (RGD) (Bachem) (PEG-RGD), and 3) PEG (diacrylate, 3.5 kDa, JenKem)-GelMA (P-G) hydrogels (n=6). All tissues were fabricated by mixing 2×10^5^ cells with the respective hydrogel solution supplemented with 0.05% Irgacure 2959 (BASF) as photoinitiator (PI) at 1:1 ratio (5μL total volume per construct was used). GelMA and PEG-RGD were synthesized following previously established protocols^51,96^ For GelMA and PEG-RGD based tissues, a final concentration of 10% hydrogel (w/v) and 0.05% PI (w/v) was used. For P-G based tissues, a final concentration of 20% PEG (3.5 kDa) (w/v), 10% GelMA (w/v), and 0.05% PI (w/v) was used. The homogenous cell distribution within the hydrogel was ensured by thorough mixing of the hydrogel solution and cell suspension by gentle pipetting. The mixture was then sandwiched between 100 μm thick spacers using a glass slide and exposed to 6.9 mW/cm^2^ UV irradiation for 30 sec. This dose of UV and photoinitiator has been shown to be safe for cell encapsulation studies^97^. The synthetic tissues were washed once with PBS (1-2 mins) and 3 times with fresh standard culture media (15 mins each) right after crosslinking to get rid of excess PI. The tissues were then maintained in standard media for 24 h prior to stress treatment.

### Mechanical characterization of hydrogels

Cell-laden hydrogels were prepared as explained in “young and aged tissue fabrication” section. Stiffness of the cell-laden hydrogels were tested using a PIUMA CHIARO nanoindenter system (Optics11, Amsterdam, The Netherlands). A colloidal probe with a spring constant of 0.48 N/m and a diameter of 98 µm was used. Three separate samples were measured for each hydrogel type and measurements were taken from 45 different locations per sample. Before testing, the sensitivity calibration of the cantilever was conducted by indenting a hard surface (i.e. a glass slide). A customized MATLAB code (The MathWorks, Inc.) was developed to determine contact points between the probe and samples and to identify Young’s moduli of the samples using the Hertz contact model:

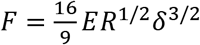

where F is applied force, δ is indentation depth, R is the radius of the colloidal probe, and E is Young’s modulus of the samples. The samples were assumed to be incompressible (i.e. Poisson’s ratio of 0.5). Statistics were performed using one-way analysis of variance (ANOVA) followed by Tukey’s multiple comparisons test, with statistical significance reported at a 95% confidence level (p < 0.05).

### Oxidative stress and epoxide hydrolase inhibitor treatment

Three different stress conditions were used: 1) via incubating the tissues in 0.2 mM H_2_O_2_ supplemented media (5% CO_2_, 37°C) for 16 h, followed by an incubation with normoxic media for 2 h. 2) via incubating the cells in an O_2_ controlled incubator at 1% O_2_ (5% CO_2_, 37°C) for 48 h in media that was pre-incubated in 1% O_2_, followed by 24 h incubation in 21% O_2_ (5% CO_2_, 37°C), and 3) via incubating the cells in an O_2_ controlled incubator at 1% O_2_ (5% CO_2_, 37°C) for 48 h in media that was pre-incubated in 1% O_2_, followed by 2 h incubation in normal conditions (21% O_2_, 5% CO_2_, 37°C). As control, tissues maintained under normoxic conditions (21% O_2_, 5% CO_2_, 37°C) for the respective treatment duration were used.

To test the effect of EHI on young and aged tissues, 0.1 mg/mL (AUDA, Sigma Aldrich) was added to culture media right before starting 48 h 1% O_2_ treatment (EHI+). This was followed by 2 h incubation under normal oxygen conditions in media supplemented with 0.1 mg/mL AUDA. The first set of control tissues used were exposed to the same hypoxia + normoxia treatment without AUDA supplement (EHI-). The second set of control tissues were maintained under normoxia conditions for the respective treatment duration.

### Live/dead assay, determining live cell percentages, normalized survival, and reactive oxygen species (ROS) levels

Following the stress treatment the samples were stained with Live/Dead assay (Life Technologies) following the manufacturer’s instructions. The tissues were imaged using a fluorescence microscope (Zeiss Hamamatsu ORCA flash 4.0) at 5 different positions (4 corners and 1 in the middle, covering approximately 5/9th of the entire construct). At each point the tissues were imaged through the thickness via 3D imaging through a built-in optical sectioning (Apotome, Zeiss) (with 12 µm scan intervals). The maximum projections of these scans were then analyzed using imageJ software to quantify live cell percentages. Using the live cell percentages, we determined the normalized cell survival in order to determine the cell death solely caused by the oxidative stress. This was done by normalizing live cell percentage of each oxidative stress treated sample to its corresponding control group (Normalized survival= (live cell% with stress/live cell% no stress control)*100).

ROS levels in tissues were determined using a ROS detection kit (Mitochondrial ROS Activity Assay Kit, Eenzyme) using manufacturer’s instructions. Briefly, the tissues were incubated with the assay agent that labels ROS products (45 min, 5% CO_2_, 37°C). Immediately after incubation the tissues were imaged using a fluorescence microscope (Zeiss Hamamatsu ORCA flash 4.0) at 5 different positions (4 corners and 1 in the middle, covering approximately 5/9th of the entire construct). The imaging through the thickness of tissues was achieved by a built-in optical sectioning (Apotome, Zeiss) (with 12 µm scan intervals). The maximum projections of these scans were then analyzed using imageJ software to determine fluorescence intensities (represented as arbitrary units) which is directly correlated with ROS levels.

### Statistical Analysis

The results are represented as average ± standard deviation. The statistical analysis was carried out using 1-way ANOVA followed by Tukey’s Multiple Comparison Test analysis. Student’s t-test was used for comparing two individual groups. All *p* values reported were two-sided, and statistical significance was defined as *p*<0.05. Sample size (n)≥3 for individual experiments.

## Acknowledgements

We thank Dr. Chad Cowan, Dr. Kiran Musunuru, Dr. Ludivine Challet-Meylan (Harvard University) for kindly providing DiPS-1016SevA cell line. We thank Dr. Athanasia Panopoulos for kindly providing Huv-iPS4F1 cell line. We thank Indiana Donor Network and Prof. Keith March (University of Florida) for kindly providing the human hearts. This study has been supported by NSF-CAREER Award # 1651385 and NSF-ECCS Award # 1611083

## Author Contributions

Aylin Acun: experimental design, collection and assembly of data, data analysis and interpretation for differentiation, molecular, functional, and age-related characterization, tissue model fabrication and characterization, manuscript writing. Trung Dung Nguyen: collection and assembly of data and data analysis for mechanical and electrophysiological characterization. Pinar Zorlutuna: design and planning of the experiments, manuscript writing.

