## Supplementary Materials for "An aged human heart tissue model showing age-related molecular and functional deterioration resembling the native heart"

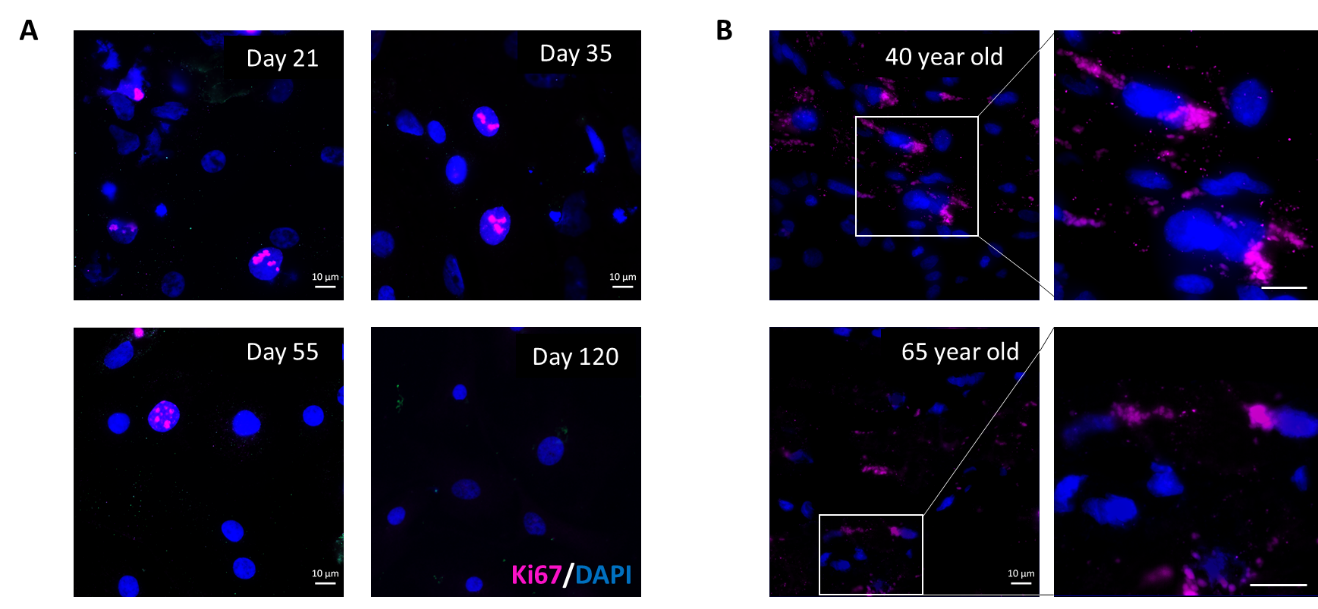

**Supplementary Figure 1.** Immunostaining against ki67 of (A) E-iCMs at different culture ages and (B) 40-year-old and 65-year-old human left ventricle sections. No nuclear localization of ki67 was detected in human left ventricle sections (Scale bars=10 μm)

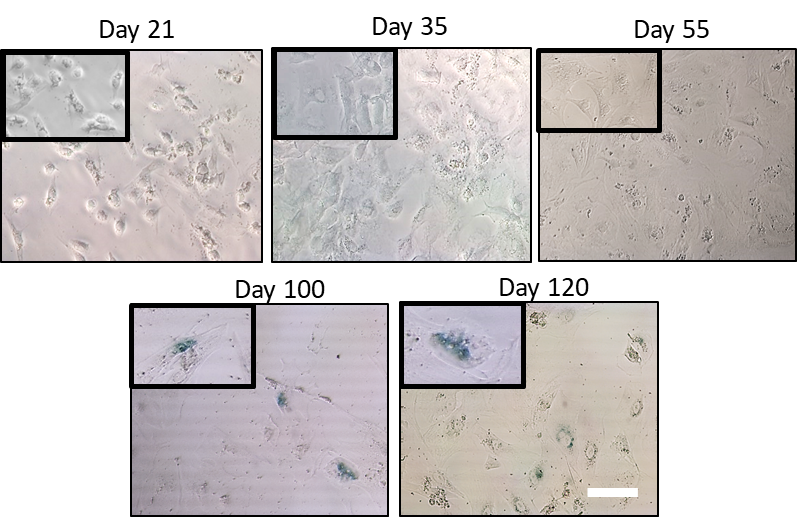

**Supplementary Figure 2.** The senescence associated β-galactosidase assay images of E-iCMs at different days of culture (blue staining indicates senescence) (Scale bar=100 μm).

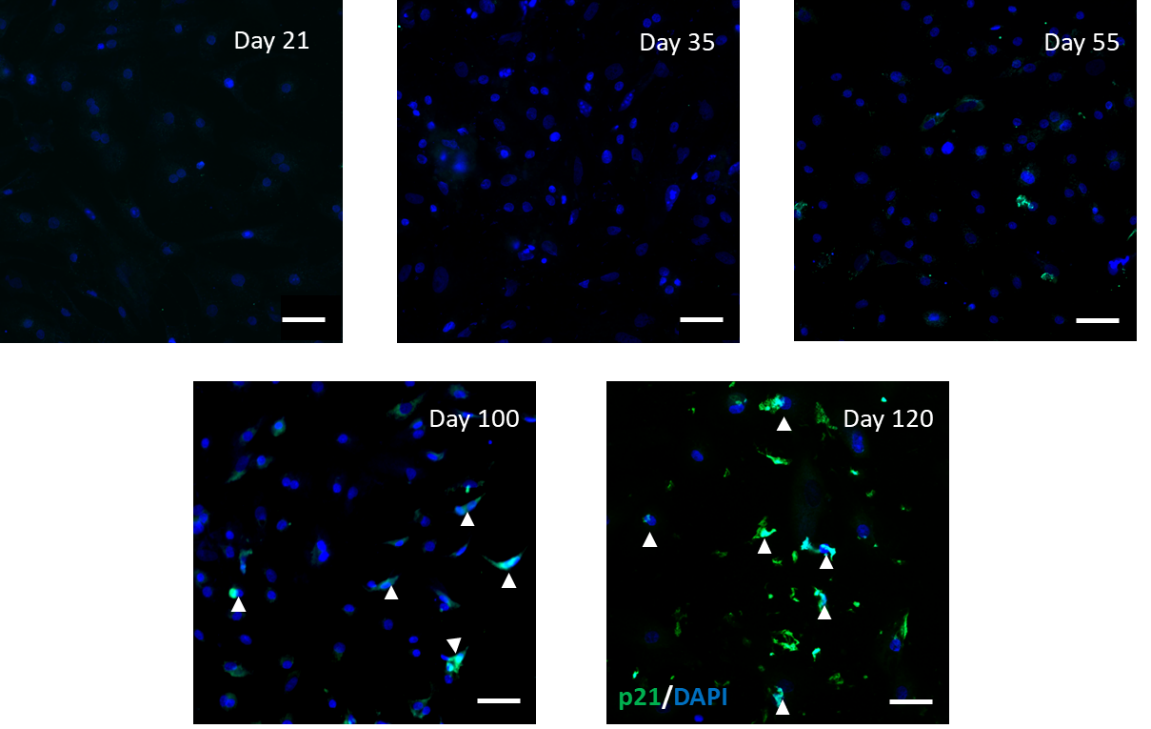

**Supplementary Figure 3.** Immunostaining of E-iCMs against p21 at different days of culture. (Scale bar=50 µm)

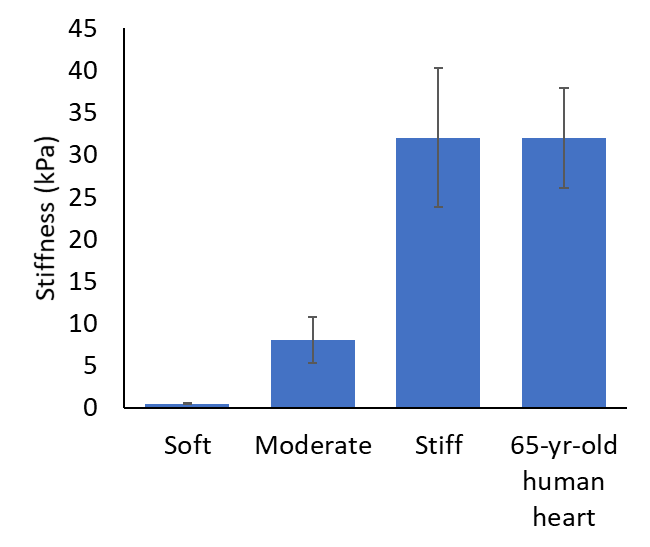

**Supplementary Figure 4.** Stiffness (kPa) of the 3 hydrogel compositions used as well as 70-year-old human heart.

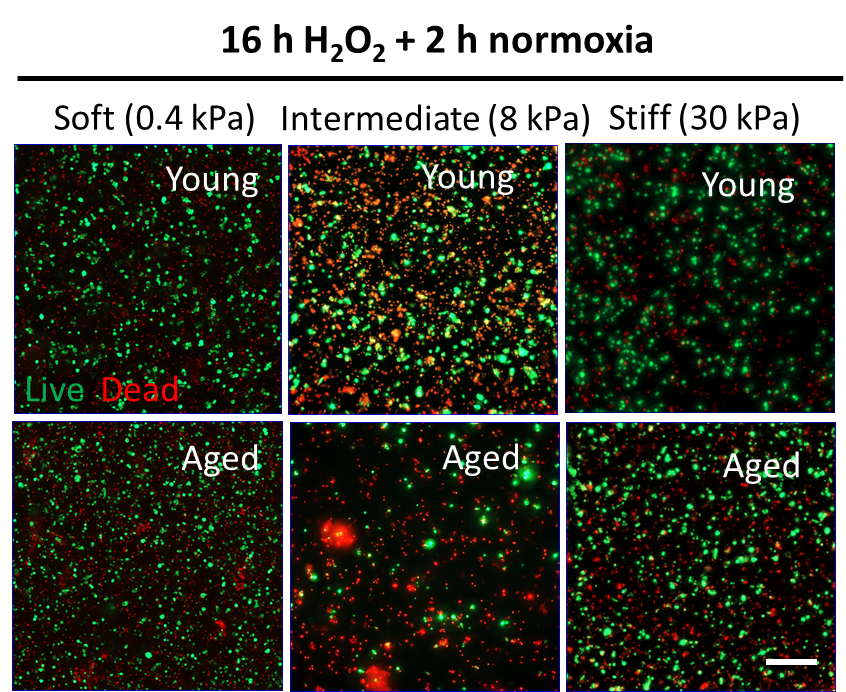

**Supplementary Figure 5.** Live/dead assay images of E-iCM soft, intermediate, and stiff young and aged tissues after exposure to 16 h H_2_O_2_ + 2 h normoxia. (Scale bar=100 μm)

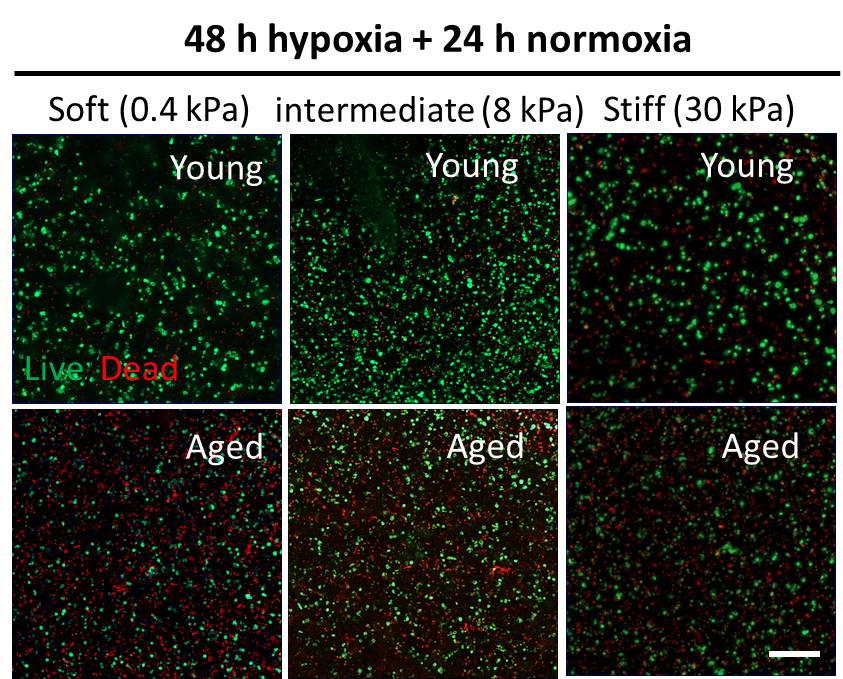

**Supplementary Figure 6.** Live/dead assay images of E-iCM soft, intermediate, and stiff young and aged tissues after exposure to 48 h hypoxia + 24 h normoxia. (Scale bar=100 μm)

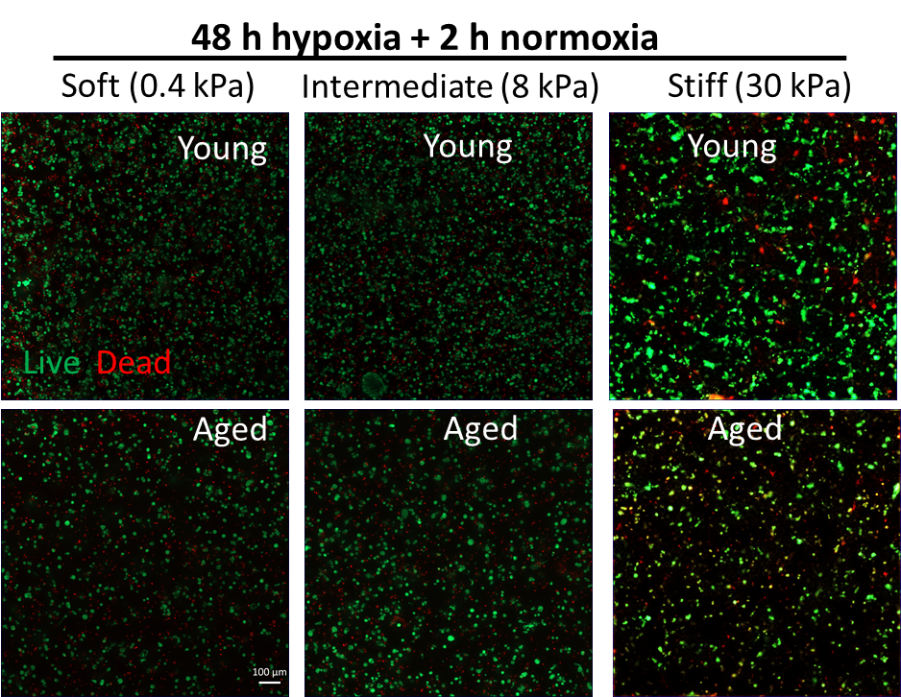

**Supplementary Figure 7.** Live/dead assay images of E-iCM soft, intermediate, and stiff young and aged tissues after exposure to 48 h hypoxia +2 h normoxia. (Scale bar=100 μm)

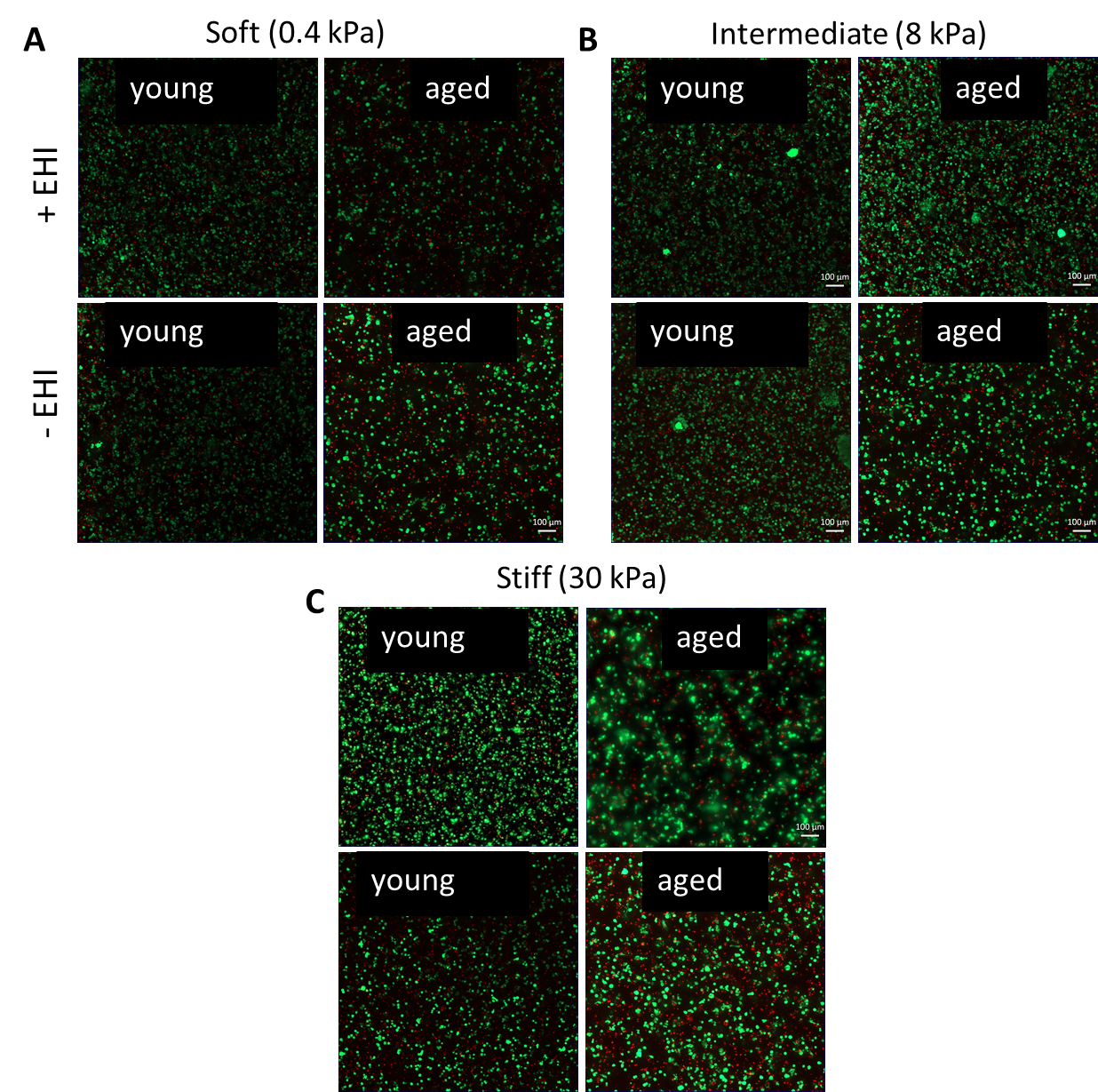

**Supplementary Figure 8.** Live/dead assay images of (A) soft, (B) intermediate, and (C) stiff young and aged E-iCM tissues with and without EHI (+EHI and –EHI, respectively). (Scale bar=100 μm)

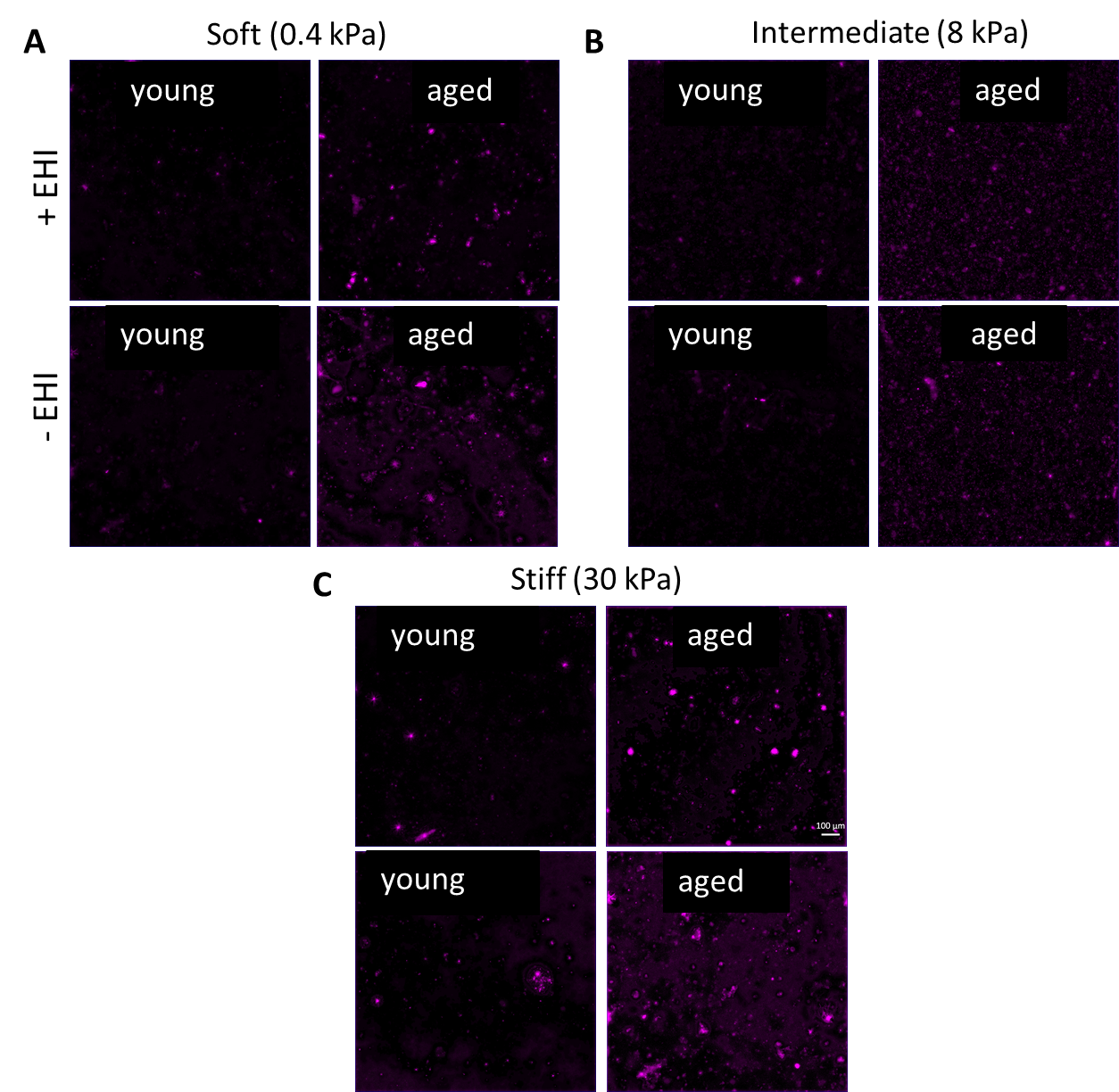

**Supplementary Figure 9.** ROS assay images of (A) soft, (B) intermediate, and (C) stiff young and aged E-iCM tissues with and without EHI (+EHI and –EHI, respectively). (Scale bar=100 μm)

| **Age** | **Sex** | **Cause of Death**  **Supplementary Table 1.** Patient information | **Other Diseases** | **Time from collection to fixation*** |
| --- | --- | --- | --- | --- |
| **40** | Male | Stroke | Hypertension | 2 h |
| **65** | Female | Stroke | Type II Diabetes | 4 h |

*The hearts were kept in organ preservation solution at 4°C**.**

**Supplementary Table 2.** List of qRT-PCR primers

| **Gene** | **Product** | **Source (Biorad)** |
| --- | --- | --- |
| GAPDH | Glyceraldehyde 3-phosphate dehydrogenase | qHsaCED0038674 |
| S100A1 | S100 Ca_2+_ binding protein A1 | qHsaCID0037853 |
| MYOM3 | Myomesin 3 | qHsaCID0018456 |
| CASQ2 | Calsequestrin-2 | qHsaCID0018297 |
| COX6A2 | Cytochrome c oxidase subunit polypeptide 2 | qHsaCED0045748 |
| TNNT2 | Cardiac troponin T 2 | qHsaCID0014544 |
| MYH6 | Myosin heavy chain 6 | qHsaCID0011216 |
| NKX 2.5 | NK 2 Homeobox 5 | qHsaCED0047574 |
| MYL2 | Myosin regulatory light chain 2 | qHsaCID0012677 |
